# Dissecting human fetal cardiac repair using cardioids

**DOI:** 10.64898/2026.06.30.735236

**Authors:** Lavinia Ceci Ginistrelli, Tobias Ilmer, Lena Plank, Maria Novatchkova, Abhijeet Krishna, Enikő Lázár, Raphaël Mauron, Stefan H. Geyer, Lokesh Pimpale, Valeria V. Orlova, Katie McDole, Wolfgang J. Weninger, Sasha Mendjan

## Abstract

Human cardiac injury responses are governed by dynamic interacting processes that are difficult to resolve. Unlike adults, fetal mammalian hearts regenerate through coordinated remodeling and proliferation supported by a pro-regenerative immune environment, extracellular matrix (ECM), and immature cardiomyocytes, including trabecular subtypes. Here, we establish a modular human cardioid injury platform to dissect these interactions. We show that anti-inflammatory macrophages selectively migrate to the injury, clear debris, and promote ECM remodeling, whereas inflammatory macrophages suppress cardiomyocyte proliferation. Synergistic FGF2-NRG1 signaling induces trabecular identity and morphology in a hyaluronan-dependent manner, conferring enhanced injury repair, characterized by cytoskeletal remodeling and cardiomyocyte proliferation mediated by YAP and WNT signaling. Exogenous YAP, but not WNT, is sufficient to promote repair in non-trabecular cardioids. These findings uncover coordinated immune-ECM-cardiomyocyte interactions governing human fetal regenerative competence and mechanistically resolve remodeling and proliferative components of cardiac repair.

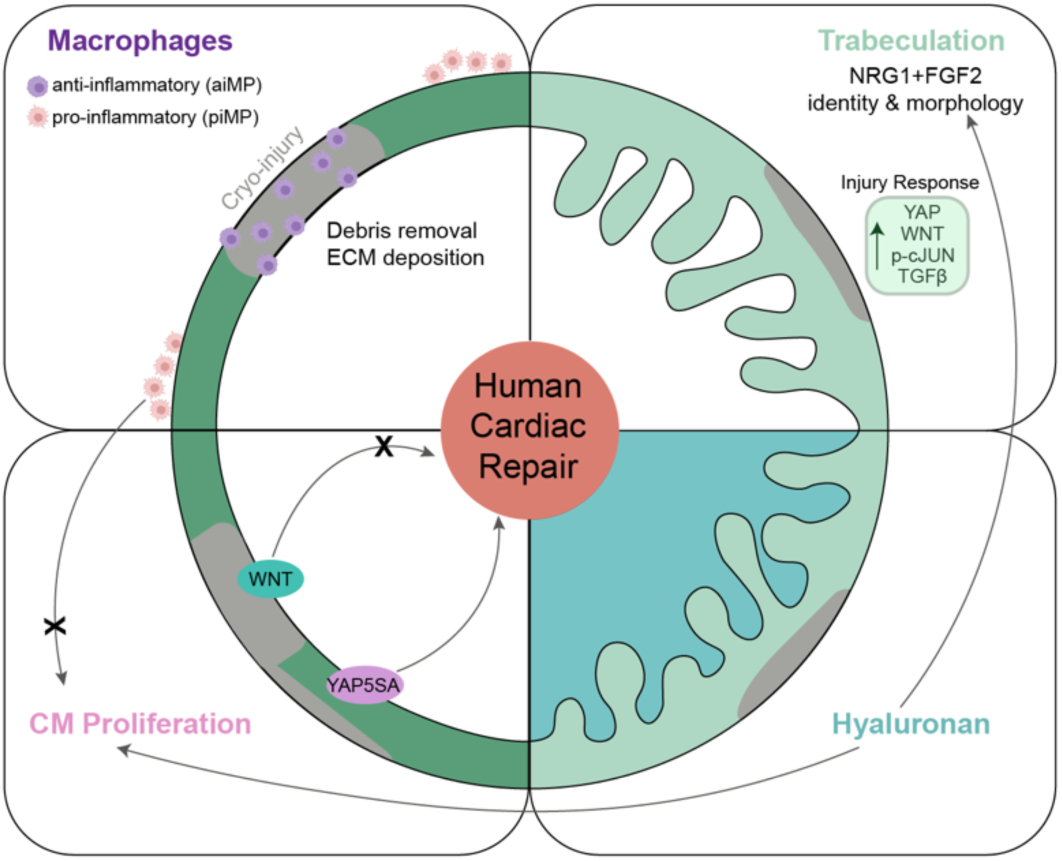

## INTRODUCTION

Loss of cardiomyocytes (CMs) following cardiac injury is a leading cause of mortality worldwide. In contrast to adult humans, newborns display a transient regenerative capacity, a phenomenon also observed in other mammals, in which fetal or neonatal heart injury can stimulate CM proliferation and tissue repair^1–3^. This regenerative competence is influenced by multiple developmental features, including an immature immune response, increased plasticity of CMs and fibroblasts, and a distinct extracellular matrix (ECM) composition. Fetal immune responses are typically less inflammatory^4^, favoring repair over fibrotic scar formation^5^. Early-stage CMs exhibit enhanced capacity to re-enter the cell cycle, and specific developmental CM subtypes, including trabecular CMs, have been implicated in regenerative responses in zebrafish and neonatal mouse models^6,7^. In addition, the embryonic and fetal cardiac ECM differs significantly from the adult, with components such as hyaluronic acid (HA) playing key roles in cardiac morphogenesis and injury responses across multiple tissues^8–13^. Despite these insights from model organisms, the extent to which these mechanisms operate in human cardiac tissue remains poorly understood.

Cardiac injury responses involve coordinated, multi-step processes integrating immune signaling, metabolic adaptation, and structural remodeling. Following injury, ECM components, including HA, contribute to tissue swelling and mechanical stabilization of the injury site^9,14^. Immune cells such as macrophages subsequently infiltrate damaged tissue to clear debris and modulate inflammatory signaling^15^. Macrophages further influence fibroblast activation states, thereby shaping whether the injury environment promotes fibrotic scar formation or tissue repair^16,17^. In non-mammalian regenerative species and neonatal mammals, pro-repair environments are associated with metabolic shifts favoring glycolysis over fatty acid oxidation and activation of signaling pathways including NRG1, YAP, and WNT, which promote CM dedifferentiation and proliferation^18–23^. However, the complexity and temporal coordination of these processes limit mechanistic understanding in human cardiac tissue, highlighting the need for the development and deployment of complementary reductionist human model systems.

Current *in vitro* models of cardiac injury do not recapitulate regenerative repair^24–28^. We previously demonstrated that human chamber-like cardiac organoids (cardioids) exhibit an intrinsic injury response characterized by fibroblast-derived collagen accumulation at sites of damage. Because cardioids form in the absence of exogenous ECM scaffolds^24^, they enable investigation of endogenous injury responses in a controlled context. Building on this platform, we established a modular regenerative cardioid injury model to systematically interrogate early phases of the fetal-like human cardiac injury response. Using this approach, we examined how macrophage recruitment, trabecular cardiac identity, and ECM composition interact to regulate cardiac remodeling and CM proliferative competence following injury.

## RESULTS

### Anti-inflammatory macrophages migrate to the injury site and contribute to ECM remodeling

Recognizing the crucial regulatory roles of macrophages in injury response^5,29,30^, we sought to integrate macrophages into the cardioid platform. To this end, we adapted an established human pluripotent stem cell (hPSC)-derived monocyte differentiation protocol^31^ that mirrors the stages of primitive hematopoietic monocyte development within the yolk sac. We first assessed differentiation efficiency and population purity by CD14+ FACS analysis and confirmed upregulation of monocyte markers (CD163, CD36, and CD45) by bulk RNA-seq analysis (Fig. S1A-B). We then confirmed that exposure to cytokines IL-4 or LPS/IFN-γ polarized CCR2- yolk sac-derived macrophages into anti-inflammatory (aiMP) or pro-inflammatory (piMP) subtypes (Fig. S1C). Specifically, bulk RNA-seq, HCR, and immunostaining validated upregulation of anti-inflammatory markers in aiMPs (MRC1, CCL13, and CD209) and upregulation of pro-inflammatory markers in piMPs (CXCL3, IL1β, and CD38) (Fig. S1D-F’). As expected, the continuous presence of the respective cytokines was required for maintaining macrophage subtype identity (Fig. S1G).

Subsequently, we co-cultured left ventricular (LV) cardioids with either aiMPs or piMPs and assessed their migration into cardioids via light-sheet live imaging and macrophage marker CD68 immunostaining at 1, 3 and 7 days post co-culture (dpc) (Fig. 1A-B’). Interestingly, the migration behavior between aiMPs and piMPs was quite distinct: aiMPs migrated faster and extensively inside cardioids as single cells (Fig. 1B-B’, Video S1), while piMPs mainly engulfed the cardioids and remained on the surface (Fig. 1B). Consistent with loss of subtype identity, removal of polarizing factors reduced migration, as evidenced by fewer CD68+ cells inside the cardioids (Fig. S1H-I). Taken together, these data support that our model system enables the dissection of distinct migratory behavior of different macrophage subtypes.

**Figure 1:**
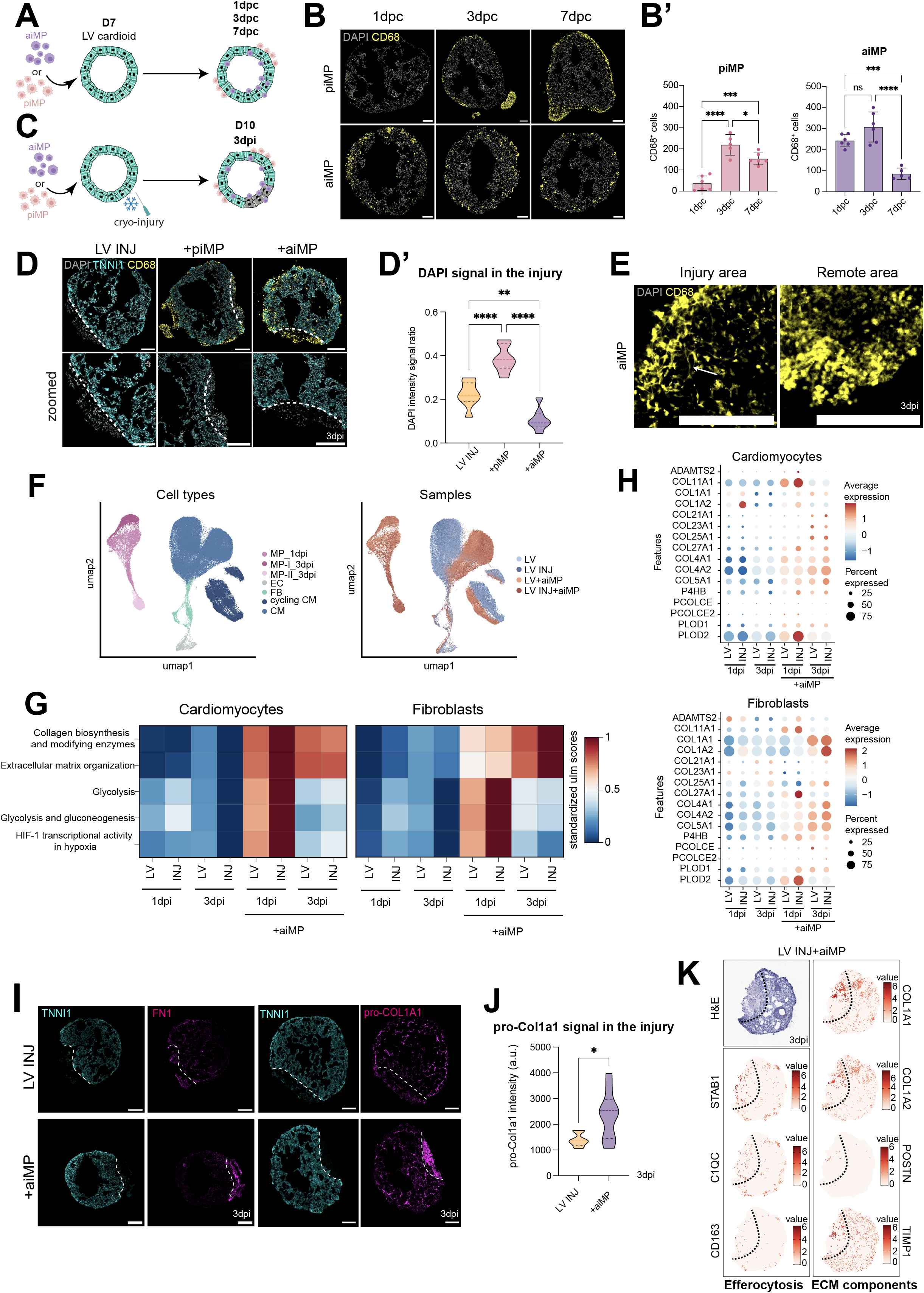
Macrophages migrate to the injury site and contribute to ECM remodeling. **A.** Schematic representation of left ventricle-like (LV) cardioids co-cultured with either pro-inflammatory macrophages (piMPs) or anti-inflammatory macrophages (aiMPs) **B.** CD68 immunostaining of LV cardioids sections co-cultured with either piMPs or aiMPs at 1-3-7 days post co-culture (dpc) with quantification (B’) (N=2, n=2-3) **C.** Schematic representation of LV cardioid cryo-injury and co-culture with either aiMPs or piMPs **D.** Cryosections of injured LV cardioids alone or co-cultured with either piMPs or aiMPs immunostained for DAPI and CD68 at 3 days post-injury (dpi). **D’.** DAPI intensity signal quantification of respective immunostaining; DAPI intensity signal in the injured area is normalized to the DAPI intensity signal in the rest of the cardioid (N=2, n=4) **E.** Representative sections of CD68 immunostaining whole-mount imaging in cryo-injured LV cardioids co-cultured with aiMPs at 3dpi. White arrow indicates CD68+ projections. **F.** UMAP representation of scRNA-seq data showing distinct groups organized by cell type or sample origin **G.** Heatmap illustrating aiMP effect in LV cardioids, showing the enrichment scores of Bioplanet terms overrepresented in genes differentially expressed between +aiMPs versus control and INJ+aiMPs vs INJ conditions in indicated cell types **H.** Dotplot showing expression of genes belonging to “Collagen biosynthesis and modifying enzymes” Bioplanet term in cardiomyocytes and fibroblasts across all conditions **I.** Fibronectin (FN) and pro-collagen1A1 (pro-COL1A1) immunostaining of cryo-injured LV cardioids with and without aiMP co-culture at 3dpi **J.** pro-COL1A1 signal intensity quantification in the injury site from whole-mount imaging of LV cardioids with or without aiMP co-culture at 3dpi (N=2, n=3-5) **K.** Spatial features plot of efferocytosis and ECM-remodeling genes in cryoinjured LV cardioids co-cultured with aiMPs at 3dpi. Scalebar: normalized expression. Scalebar is 200 µm. Dashed line highlights the injured area. Bar graphs show mean ± SD. Statistics: one-way ANOVA. *p < 0.05, **p < 0.01, ***p < 0.001, ****p < 0.0001.

We next used this platform to model the human fetal cardiac injury response using cryo-injured LV cardioids cultured in the presence of aiMPs or piMPs (Fig. 1C). Through live imaging and CD68 immunostaining analysis, we demonstrated that aiMPs robustly migrate to the injury site, characterized by loss of cardiac reporter and increased DAPI signal, and maintain their localization over time (Fig. 1D, S1J, Video S2). However, piMPs did not consistently localize at the injury site (Fig. 1D, S1K, Video S3). We found that aiMPs were functionally active at the injury site, as evidenced by decreased intensity of DAPI signal, likely reflecting removal of debris (Fig. 1D-D’). We also noted that aiMPs acquired an elongated morphology, distinct from a rounded morphology elsewhere in the cardioid, possibly indicating different phenotypic states^32^ (Fig. 1E).

To identify candidate molecular programs activated during aiMP-CM interactions upon injury, we performed single-cell RNA-sequencing (scRNA-seq) on LV cardioids cultured in the presence or absence of aiMPs, with and without cryo-injury, at 1 and 3 days post-injury (dpi). Unsupervised clustering identified the major cardiac cell populations, including cycling and non-cycling CMs, fibroblasts (FB), macrophages (MP, MP-I, MP-II), and endothelial cells (EC) (Fig. 1F), confirmed by canonical marker gene expression (Fig. S1L). Differential expression analysis showed that aiMP co-culture strongly altered CM transcriptional identity, particularly at the early time point (1 dpi) (Fig. 1G). CMs exposed to aiMPs upregulated gene programs associated with glycolysis and glucose metabolism while downregulating oxidative phosphorylation pathways (Fig. 1G, Fig. S1M-N), consistent with *in vivo* transient metabolic reprogramming upon injury^18,33^. In parallel, the presence of aiMPs induced extracellular matrix (ECM)-related genes, including collagen biosynthesis and modifying enzymes such as COL1A1 and PLOD1/2 in both CMs and fibroblasts (Fig. 1G–H). These effects were more pronounced following cryo-injury, consistent with activation of early fibrotic remodeling programs (Fig. 1G–H). Consistent with these transcriptional changes, immunostaining confirmed increased deposition of ECM proteins, including fibronectin (FN) and pro-COL1A1, specifically at injury site in cardioids co-cultured with aiMPs (Fig. 1I–J). This effect was independent of cytokine exposure (Fig. S1O), supporting a direct role of aiMPs in promoting fibrotic ECM remodeling and consistent with *in vivo* observations that macrophages regulate collagen deposition following injury^15,34^.

To further resolve the spatial distribution of cardiomyocytes, fibroblasts and the two populations of macrophages at 3dpi, we performed Visium HD spatial transcriptomics analysis on cryoinjured LV cardioids co-cultured with aiMPs. scRNA-seq cluster deconvolution showed preferential localization of fibroblasts and MP-I at the injury site (Fig. S1P). MP-I exhibited enriched expression of canonical anti-inflammatory markers and efferocytosis-associated genes, suggesting that these cells resemble anti-inflammatory macrophages involved in apoptotic cell clearance (Fig. S1Q-R). Localized expression of efferocytosis-associated genes, including STAB1 and C1Q genes, within the injury site, supports macrophage activation in debris clearance (Fig. 1K). Spatial feature plots displaying ECM components confirmed enriched expression of collagens, including COL1A1 and COL1A2, and ECM-remodeling genes, such as POSTN and TIMP1, at the injury site (Fig. 1K).

Together, these results show that incorporation of defined macrophage subtypes into the cardioid injury platform reveals that aiMPs preferentially localize to the injury site, promote debris clearance, and stimulate ECM deposition, thereby coordinating both metabolic and reparative responses in cardiomyocytes and fibroblasts.

### Synergistic NRG1 and FGF2 signaling induces trabecular identity, morphology, and injury remodeling

The cardioid–macrophage co-culture system enabled the dissection of macrophage subtype-specific injury responses. Notably, during development, macrophages first colonize the human heart at the trabeculation stage, and trabecular myocardium plays a key role in cardiac repair across species, including humans^7,35,36^. Thus, we reasoned that further developing our cardioid model to recapitulate the trabeculation stage of heart development would provide a powerful tool to investigate the complex contribution of trabecular myocardium to cardiac injury and repair in humans.

Trabecular CMs arise from early ventricular CMs and are defined by distinct marker gene expression (NPPA+, IRX3+, BMP10+, GJA5+), morphology, and dependence on ECM components such as HA (which itself has a vital role in injury)^8,37,38^. Trabeculation is mediated by endocardial-myocardial crosstalk involving neuregulin (NRG1)-ERBB signaling^39–41^. To model this crosstalk, we treated cardioids with NRG1 alone, FGF2 alone, or NRG1 combined with FGF2 (NF). While all treatments increased cardioid size (Fig. 2A-A’), only NF treatment robustly induced expression of all canonical trabecular markers, including BMP10, at 3 days post-treatment, as confirmed by RNA-seq and HCR (Fig. 2B-C’). We therefore refer to these cardioids as trabecular left ventricular (TLV) cardioids.

**Figure 2:**
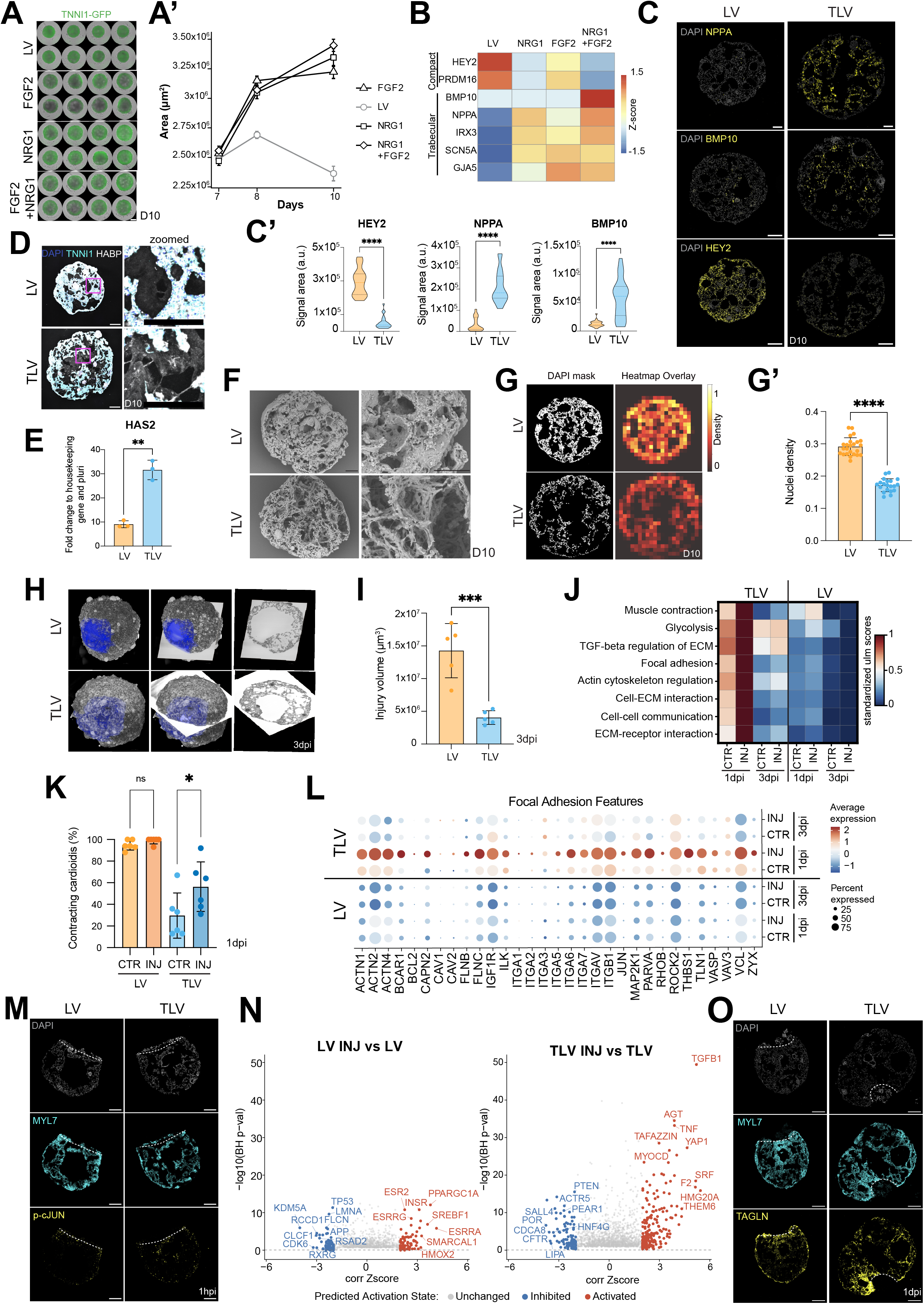
Synergistic NRG1 and FGF2 signaling induces trabecular identity, morphology, and injury remodeling. **A.** Widefield images of TNNI1-GFP-derived LV cardioids treated with NRG1, FGF2 and NRG1+FGF2 at 3 days post treatment (D10). Scale bar is 500 µm **A’.** Size quantification of the respective conditions at D7-D8-D10 (N=3, n=8) **B.** RNA-seq expression heatmap of trabecular and compact gene markers in indicated conditions at D10 **C.** NPPA, BMP10 and HEY2 HCR staining and quantification (C’) of LV and TLV cardioid sections at D10 (N=3, n=4-8) **D.** HABP immunostaining of LV and TLV cardioid cryosections at D10 **E.** RT-qPCR of HAS2 in LV and TLV cardioids at D10 (N=3) **F.** Scanning Election Microscopy (SEM) images of LV and TLV cardioids at D10 **G-G’.** Nuclei density quantification of LV and TLV cardioids DAPI-stained sections (N=3, n=8-16) **H.** Representative images of High Resolution Episcopic Microscopy (HREM) of LV and TLV cryo-injured cardioids at 3dpi (D10); injury volume is colored in blue **I.** Injury volume quantification from whole mount imaging of cryo-injured LV and TLV cardioids at 3dpi (N=3, n=1-2) **J.** Heatmap illustrating INJ effect in LV and TLV cardioids, showing the enrichment scores of Bioplanet terms overrepresented in genes differentially expressed between INJ versus CTR CMs, with analyses performed independently for LV and TLV at both timepoints **K.** Percentage of contracting cardioids in the respective conditions at 1dpi (N=6, n=8-16) **L.** Dotplot showing expression of genes belonging to “Focal adhesion” Bioplanet term in cardiomyocytes in the indicated conditions **M.** Phospho-cJUN immunostaining in cryoinjured MYL7-GFP-derived LV and TLV cardioids at 1hpi **N.** Volcano plot of IPA upstream analysis highlighting predicted upstream regulators associated with genes differentially expressed in LV INJ vs LV and TLV INJ vs TLV ctr at 1dpi **O.** TAGLN immunostaining in cryoinjured MYL7-GFP-derived LV and TLV cardioids at 1dpi. Scalebar is 200 µm, except where specified. Dashed line highlights the injured area. Bar graphs show mean ± SD. Statistics: Student’s t-test. *p < 0.05, **p < 0.01, ***p < 0.001, ****p < 0.0001.

TLV cardioids exhibited reduced CM proliferation, as shown by TNNT2:FUCCI FACS analysis (Fig. S2A), suggesting that increased cardioid size reflects altered tissue architecture rather than increased cell division.

*In vivo*, trabeculation critically depends on HAS2-mediated HA production^8,42^. Consistent with this, NF-treatment induced upregulation of HAS2 levels and increased HABP signal (Fig. 2D-E), indicating increased HA deposition. This supports the conclusion that NF signaling promotes trabecular morphogenesis through ECM remodeling.

Scanning electron microscopy revealed ridge-like CM structures resembling trabecular architecture (Fig. 2F), accompanied by reduced nuclear density and altered actin organization (Fig. 2G-G’, S2B). Withdrawal of NF signaling led to loss of trabecular markers (Fig. S2C), indicating that sustained NF signaling is required to maintain trabecular CM identity in the absence of endocardial cells.

Because macrophages first colonize the fetal heart during trabeculation, we next investigated whether CM identity influences macrophage localization. Co-culture of aiMPs with control LV or TLV cardioids revealed reduced macrophage infiltration into trabecular-like structures (Fig. S2D-D’), consistent with *in vivo* observations that CCR2- macrophages preferentially localize outside of the trabecular compartment^43^. Altogether, these data establish TLV cardioids as a model of trabeculation-stage myocardium suitable for dissecting subtype-specific immune interactions during injury.

Given the critical role of trabecular signaling and structures in cardiac remodeling and regeneration, we next utilized trabecular cardioids to further dissect the effects of injury on different CM subtypes. Cryo-injured TLV cardioids exhibited markedly reduced injury areas compared to LV cardioids, with minimal accumulation of debris-like structures (Video S4, Fig. S2E). High-Resolution Episcopic Microscopy (HREM), which enables 3D reconstruction and volumetric analysis of entire cardiac organoids, confirmed a reduction in injury volume in TLV cardioids relative to LV cardioids (Fig. 2H, S2F), further supported by whole-mount volumetric quantification showing an approximately threefold decrease in injured tissue (Fig. 2I). These results indicate that trabecular cardioids exhibit enhanced injury remodeling capacity, suggesting CM subtype-specific differences in regenerative potential.

To further define the molecular basis of these differences, we performed scRNA-seq on injured and uninjured LV and TLV cardioids at 1 and 3 dpi, with and without macrophage co-culture. Unsupervised clustering identified cardiomyocytes, cycling cardiomyocytes, fibroblasts, macrophages, and endothelial cells (Fig. S2G-H). Interestingly, despite reduced aiMP infiltration in TLV cardioids, macrophage co-culture induced upregulation of glycolysis-related genes in TLV CMs (Fig. S2I). These findings suggest that aiMPs can promote metabolic reprogramming in both LV and TLV cardioids.

To resolve the spatial distribution of TLV cardiomyocytes and macrophages, we performed Visium HD spatial transcriptomics analysis on cryoinjured TLV cardioids co-cultured with aiMPs at 3dpi. scRNAseq deconvolution showed that, similar to LV cardioids, MP- I, characterized by enrichment of anti-inflammatory and efferocytosis markers (Fig. S2J-K), preferentially localizes at the injury site (Fig. S2L). Spatial feature plots confirmed localization of efferocytosis and ECM components genes, including STAB1, CD163, COL1A1 and POSTN, at the injury site (Fig. S2M). These findings suggest that aiMPs migrate and are functionally active at the injury site also in TLV cardioids.

scRNA-seq differential expression analysis of TLV cardioids in the absence of macrophages revealed that trabecular CMs exhibit a stronger injury-induced transcriptional response than LV CMs (Fig. 2J). At 1dpi, injured TLV CMs showed marked enrichment of genes and Gene Ontology (GO) terms associated with cardiac remodeling, including muscle contraction (MYH7), glycolysis, focal adhesion components (integrins, actin, collagens, integrin-linked kinase), ECM organisation, cell-ECM interactions, and actin cytoskeleton remodeling (Fig. 2J, 2L). These transcriptional signatures are consistent with *in vivo* studies linking chamber mechanobiology genes, such as NPPA, NPPB, and ANKRD1, to injury response (Fig. S2N).^44–47^ Functional analysis supported these findings: contraction measurements revealed a higher proportion of contracting TLV cardioids following cryoinjury relative to uninjured control at 1 dpi (Fig. 2K), indicating enhanced functional adaptation to injury. Among focal adhesion-associated genes, JUN was strongly induced upon injury at 1dpi (Fig. 2L), suggesting activation of AP-1-mediated stress signaling. AP-1 is known to rapidly activate injury-responsive loci associated with tissue remodeling and regeneration^48^, suggesting that trabecular NF signaling sensitizes cardiomyocytes to stress-induced remodeling programs. Immunostaining for phospho-cJUN one hour after injury confirmed increased pathway activation in injured TLV CMs (Fig. 2M, S2O), consistent with rapid engagement of injury-responsive transcriptional programs. Ingenuity Pathway Analysis (IPA), further identified of TGFβ- and YAP-associated signaling pathways (Fig. 2N), prominent regulators of cardiac remodeling and regenerative responses^20,49,50^. Immunostaining for TAGLN, a known TGFβ target involved in cardiomyocytes injury remodeling^51^, confirmed upregulation of TAGLN in TLV cardioids compared to control LV and localization of TAGLN+ CM around the injury site at 1dpi (Fig. 2O, S2P). In summary, we demonstrate that combined NRG1 and FGF2 signaling is both necessary and sufficient to induce trabecular cardiomyocyte identity and morphology in cardioids. Using this platform, we show that macrophage recruitment is influenced not only by macrophage subtype but also by cardiomyocyte identity and that TLV cardioids display a distinct early injury-response characterized by enhanced cytoskeletal reorganization, metabolic remodeling, contractile adaptation, and rapid AP-1 pathway activation.

### Hyaluronic acid (HA) regulates trabecular identity and injury remodeling

*In vivo*, the fetal cardiac ECM, particularly its major component HA, plays a critical role in trabeculae formation and contributes to both pathological remodeling, such as myocardial infarction, and regenerative wound-healing responses^9,52^. We recently showed that the mechanobiological properties of HA are essential for cardioid morphogenesis, CM specification, and contractile capability, enabling functional dissection of ECM contributions that are otherwise difficult to resolve^53^. Given the increased HAS2 expression observed in TLV cardioids (Fig. 2E), we hypothesized that trabecular identity and its associated injury response are regulated by HA-dependent mechanisms.

To test this hypothesis, we enzymatically degraded HA using hyaluronidase (HAase). HA depletion resulted in a dose-dependent reduction in both LV and TLV cardioid size, accompanied by cavity collapse and increased cell density (Fig. 3A-C’’, S3A-C’). In TLV cardioids, HA degradation also reduced expression of trabecular markers, suggesting that ECM composition directly influences CM identity and that HA is required for trabecular specification (Fig. 3D-E’, S3D).

**Figure 3:**
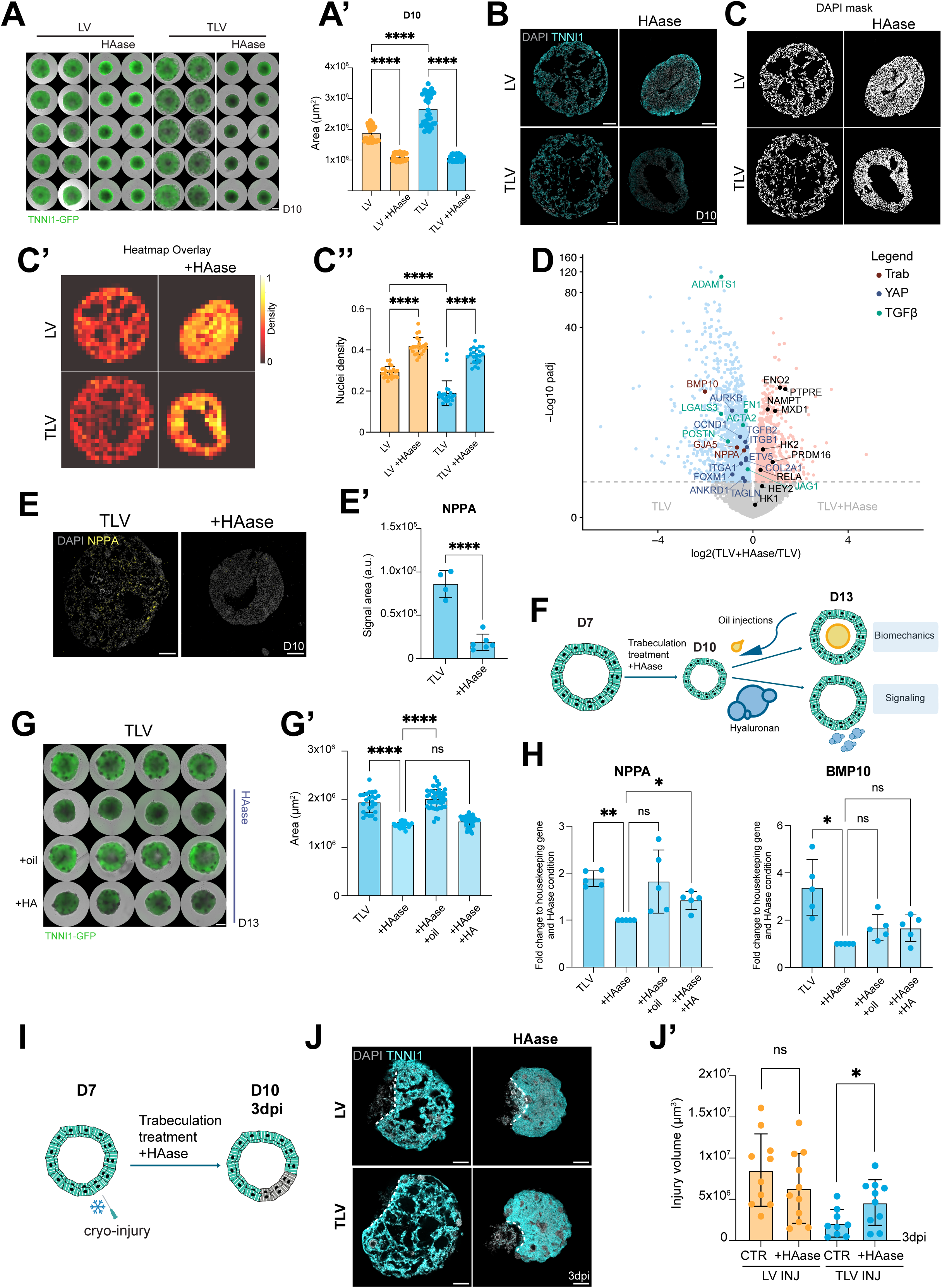
Hyaluronic acid regulates trabecular identity and injury remodeling. **A.** Widefield images of TNNI1:GFP-hPSC-derived LV and TLV cardioids untreated and treated with hyaluronidase (HAase) at 3 days post treatment (D10) with size quantification (A’) (N=3, n=9-11). Scale bar is 500µm **B.** Cryosections of TNNI1:GFP-hPSC-derived LV and TLV cardioids untreated and treated with HAase **C-C’’.** Nuclei density quantification of indicated conditions at D10 with representative DAPI and heatmap masks (N=3; n=6-8). **D.** RNA-seq volcano plot showing differentially expressed genes in indicated conditions. NS=Not Significant; UP=Upregulated genes; DOWN=Downregulated genes; Trab=Trabecular marker genes; YAP=YAP target genes; TGFβ=TGFβ target genes. **E.** NPPA HCR staining of TLV cardioids sections treated with HAase at D10 (N=2, n=4-6) and quantification (E’)**. F.** Schematic representation of rescue experiment with oil injection and HA added to the media **G.** Widefield images of TNNI1:GFP-hPSC-derived TLV cardioids treated as indicated with size quantification (G’) at D13 (N=3, n=8-20). Scale bar is 500 µm. **H.** RT-qPCR of NPPA and BMP10 in indicated conditions at D13 (N=5, n=8). **I.** Schematic representation of cryo-injured and trabeculation/HAase-treated cardioids **J.** Representative sections of whole mount images of injured TNNI1:GFP-hPSC-derived LV and TLV cardioids in indicated conditions at 3dpi. Dashed line highlights the injured area. **J’** Injury volume quantification from whole-mount images of indicated conditions at 3dpi (D10) (N=3, n=3-4). Statistics: Student’s t-test Scalebar is 200 µm, except where specified. Bar graphs show mean ± SD. Statistics: one-way ANOVA, except where specified. *p < 0.05, **p < 0.01, ***p < 0.001, ****p < 0.0001.

To determine whether HA is required for trabeculae initiation, we treated cardioids with HAase (D5-D7) prior to NF treatment (Fig. S3E). Early HA depletion reduced cardioid size and impaired trabecular CM specification, as indicated by reduced NPPA and BMP10 expression (Fig. S3F-G), suggesting that HA might be necessary for initiation of trabecular identity. In contrast, HA degradation (D10-D13) after trabecular specification reduced cardioid size without significantly altering expression of trabecular markers (Fig. S3H-J), suggesting that HA is required for establishment but not maintenance of CM subtype identity. Collectively, these results define stage-specific roles of HA during trabecular morphogenesis.

To distinguish mechanical and signaling contributions of HA to trabecular cardiomyocyte identity, we next tested whether trabecular features could be rescued following HA depletion (Fig. 3F). Injection of inert oil into HA-depleted TLV cardioids restored organoid size (Fig. 3G-G’) and partially rescued expression of key trabecular markers, including NPPA and BMP10 (Fig. 3H), indicating that mechanical outward expansion contributes to trabecular morphogenesis. In contrast, supplementation with exogenous HA following enzymatic degradation did not restore cardioid size but partially recovered expression of main trabecular markers, NPPA and BMP10 (Fig. 3G-H), supporting a distinct signaling role of HA. Together, these findings demonstrate that both the signaling and structural properties of HA contribute to trabecular CM identity and tissue morphogenesis.

We next asked whether HA contributes to the enhanced injury responsiveness observed in TLV cardioids (Fig. 3I). HA degradation increased injury size in TLV cardioids at 3dpi compared to untreated control, whereas injury volume in LV cardioids was not significantly affected by HAase treatment (Fig. 3J-J’). Consistent with TLV cardioid remodeling phenotype after injury, HA degradation also altered expression of TGFβ- and YAP-associated genes, including ADAMTS1, FN1, POSTN, TGFβ2, AURKB2, integrins, and ANKRD1 (Fig. 3D), further linking HA-dependent mechanobiology to injury-responsive signaling pathways. These findings indicate that HA plays a selective role in trabecular injury remodeling and contributes to the heightened injury responsiveness of TLV cardioids.

Taken together, these results demonstrate that the cardioid platform enables reductionist dissection of key determinants of cardiac injury biology, including macrophage recruitment, trabecular-specific remodeling, and fetal ECM-mediated regulation of injury resolution.

### Integrated cardiomyocyte proliferation and remodeling is necessary for cardioid injury repair

In addition to ECM remodeling, cytoskeletal reorganisation and immune signaling, cardiac repair also requires coordinated regulation of CM proliferation. Because both tissue remodeling and CM proliferation contribute to regenerative outcomes, we systematically examined how injury, macrophage subtype, trabecular signaling, and HA mechanobiology influence CM proliferative responses.

We first tested whether the difference in TLV cardioid injury response might also be due to differences in CM proliferation. EdU labeling revealed increased proliferation in TLV CMs at 3 dpi, whereas LV cardioids showed no significant change to uninjured controls (Fig. 4A). Consistently, TNNT2:FUCCI analysis demonstrated a significant increase in cycling Clover+ CMs specifically in injured TLV cardioids (Fig. S4A). scRNA-seq analysis further confirmed an increased proportion of cycling CMs in TLV relative to LV cardioids following injury (Fig. S4B). Together, these data suggest that TLV CMs show a more proliferative response upon cryo-injury, despite exhibiting overall lower baseline proliferation levels compared to LV CMs.

**Figure 4:**
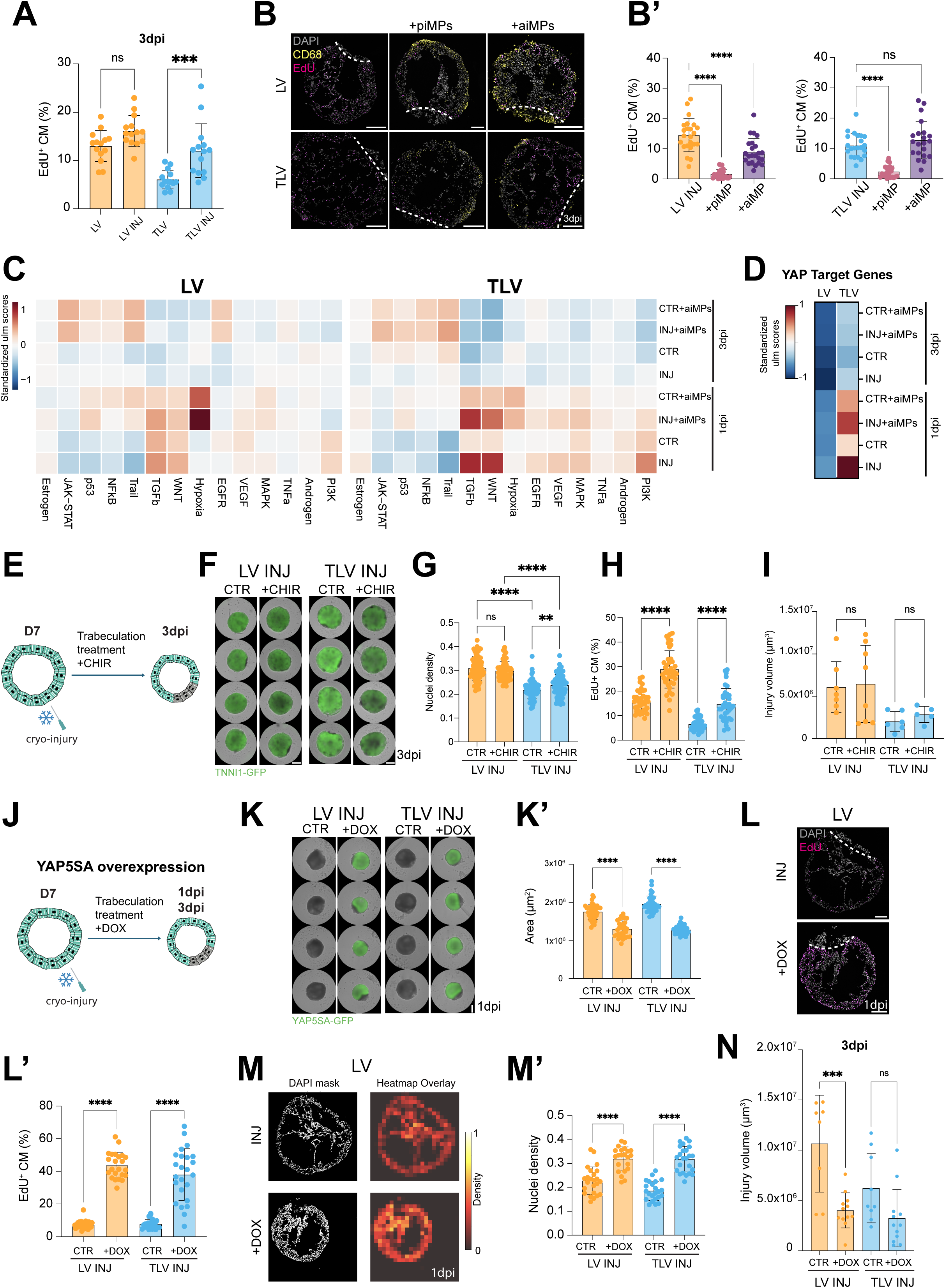
Integrated cardiomyocyte proliferation and remodeling is necessary for cardioid injury repair. **A.** Quantification of EdU+ CM in indicated conditions at 3dpi (N=3, n=4-6). **B.** Representative images of LV cardioid alone and co-cultured with either piMPs or aiMPs stained for EdU and CD68 with quantification (B’) of EdU+ CM in indicated conditions at 3dpi (N=3, n=8). **C.** Pathway activity score heatmap using PROGENy analysis in cycling cardiomyocytes in indicated conditions **D.** Heatmap showing expression of YAP-upregulated target genes identified by unbiased IPA analysis. Standardized across all shown conditions. **E.** Schematic protocol representation of cryo-injured cardioids treated with trabeculation and/or CHIR treatment at indicated timepoints **F.** Widefield images of TNNI1:GFP-hPSC-derived LV and TLV cardioids in indicated conditions at 3dpi. Scalebar is 500 µm **G.** Nuclear density quantification of indicated conditions (N=4, n=6-10) **H.** Quantification of EdU+ CM in indicated conditions at 3dpi (N=4, n=6-8) **I.** Injury volume quantification from whole-mount images of indicated conditions at 3dpi (N=2, n=3-4) **J.** Schematic protocol representation of YAP5SA:GFP-hPSC-derived cardioids treated with trabeculation and/or DOX treatment at indicated timepoints **K.** Widefield images of YAP5SA:GFP-hPSC-derived LV and TLV cardioids in indicated conditions with size quantification (K’) at 1dpi (N=3, n=12). Scale bar is 500 µm **L.** Representative images of untreated and DOX-treated LV cardioid cryosections stained for EdU and DAPI. **L’.** Quantification of EdU+ CM in indicated conditions at 1dpi (N=3, n=8). **M.** Representative heatmap mask of nuclei density of untreated and DOX-treated LV cardioids cryosections at 1dpi **M’.** Nuclei density quantification of indicated conditions at 1dpi (N=3, n=8). **N.** Injury volume quantification from whole-mount images of indicated conditions at 3dpi (N=3, n=3-4). Scalebar is 200 µm, except where specified. Dashed line highlights the injured area. Bar graphs show mean ± SD. Statistics: one-way ANOVA. *p < 0.05, **p < 0.01, ***p < 0.001, ****p < 0.0001.

Given that TLV cardioids exhibit HA-dependent injury response and that HA regulates proliferation in multiple cell types^54^, we next examined whether HA mechanobiology contributes to CM proliferative competence. Enzymatic degradation of HA significantly reduced CM proliferation in both LV and TLV cardioids at 3dpi, as measured by PHH3 staining in whole mount images (Fig. S4C), indicating that ECM composition influences mitogenic responses.

Based on prior evidence that macrophages modulate CM proliferation *in vivo*^55^, we then evaluated the effects of piMPs and aiMPs on CM proliferation following cryo-injury. EdU incorporation and TNNT2:FUCCI FACS analysis revealed that both macrophage subtypes reduced CM proliferation in LV cardioids, with a stronger inhibitory effect observed for piMPs (Fig. 4B-B’, S4D). In TLV cardioids, piMP co-culture similarly decreased CM proliferation, whereas aiMPs showed a modest but non-significant trend toward increased proliferation (Fig. 4B’). Immunostaining further indicated that reduced proliferation in the presence of piMPs was localized near macrophages (Fig. 4B), consistent with a local suppressive effect on neighboring CMs.

To investigate signaling pathways associated with the proliferative response, we performed pathway activity analysis on cycling CMs from the scRNAseq dataset (Fig. 4C). This analysis revealed pronounced activation of multiple signaling pathways in TLV CMs, particularly at 1 dpi. aiMP co-culture preferentially enhanced hypoxia-associated transcriptional programs consistent with glycolytic metabolic adaption, whereas injury alone induced upregulation of TGFβ, WNT, MAPK, and PI3K signaling in TLV CMs (Fig. 4C). Expression analysis of YAP targets confirmed strong early activation of YAP signaling targets in proliferating TLV CMs at 1 dpi, although this response was slightly blunted in the presence of aiMPs (Fig. 4D). By 3 dpi, activation of these pathways was reduced, indicating that TLV CMs mount a rapid but transient proliferative signaling response following injury (Fig. 4C-D).

We next tested whether activation of pathways identified in TLV cardioids is sufficient to promote repair in injured LV cardioids (Fig. 4E, 4J). Treatment with the WNT activator CHIR increased CM size in LV cardioids, but not in TLV cardioids (Fig. S4E). However, TLV cardioids showed increased nuclear density at 3dpi (Fig. 4G). CM proliferation was upregulated in both LV and TLV cardioids at 3 dpi (Fig. 4H, S4F). However, WNT activation did not reduce injury volume (Fig. 4I), indicating that increased proliferation alone is not sufficient to promote a pro-reparative response.

Given that TLV cardioids strongly upregulated YAP targets following injury in an HA-dependent manner (Fig. 2-3), we next tested whether YAP activation is sufficient to promote reparative responses in LV cardioids. Using a doxycycline-inducible YAP5SA construct encoding a constitutively active YAP^56^ (Fig. 4J, S4G-H), we observed increased cardiomyocyte proliferation in both LV and TLV cardioids (Fig. 4L-L’), accompanied by more compact tissue morphology characterized by reduced size and increased nuclear density at 1 dpi (Fig. 4K-K’, 4M-M’). CM proliferation returned to near baseline levels by 3 dpi (Fig. S4K), whereas the compact morphology persisted (Fig. S4I-J), indicating that transient YAP activation induces a short-lived proliferative burst. Importantly, YAP activation reduced injury volume in both LV and TLV cardioids (Fig. 4N), demonstrating that YAP promotes both proliferative and remodeling responses in injured cardiac tissue.

In conclusion, these findings suggest that trabeculation enhances responsiveness to regenerative cues through coordinated activation of mechanosensitive remodeling pathways and transient mitogenic signaling programs. Whereas WNT activation increases cardiomyocyte proliferation without improving repair, YAP activation promotes both proliferation and injury remodeling, suggesting that effective cardiac repair requires integration of structural remodeling and proliferative competence. Overall, these results demonstrate that interactions between macrophages, ECM mechanobiology, and cardiomyocyte identity converge to regulate injury responses in human cardiac tissue.

## DISCUSSION

Here, we established a human cardioid injury platform that enables systematic dissection of fetal cardiac injury response within defined ECM and cellular contexts, including distinct macrophage (pro- and anti-inflammatory) and CM (non-/ventricular and trabecular) subtypes. While the model remains inherently reductionist relative to the human fetal heart, this simplified system enables controlled reconstruction of complex regenerative processes in a stepwise manner, beginning with a minimal set of cardiac cell types. The modular design of the platform allows incorporation of additional cell types, including epicardial-derived fibroblasts, endothelial cells, and diverse immune populations, enabling mechanistic interrogation of multicellular interactions that regulate cardiac repair. Moreover, the system permits targeted perturbation of signaling pathways implicated in cardiac regeneration, including NRG, FGF, WNT, and YAP, whose cell type–specific roles and temporal dynamics across regenerative stages remain incompletely understood, particularly in human tissue. Because cardioids maintain an endogenous ECM, the platform also enables functional interrogation of matrix composition, providing experimental access to mechanobiological processes that are otherwise difficult to isolate. In addition, the system opens the gateway to in-depth biochemical analyses, including time-resolved phospho-proteomic profiling, linking molecular signaling dynamics to emergent repair phenotypes.

Application of this platform revealed several biological insights into determinants of injury responsiveness in human cardiac tissue. We show that anti-inflammatory macrophages selectively localize to sites of injury and promote fibroblast-driven extracellular matrix deposition and remodeling, similar to reparative macrophages *in vivo*^57,58^. They also transiently induce a hypoxia and glycolysis transcriptional signature, suggesting a possible shift towards a regeneration-permissive microenvironment^59^. Conversely, pro-inflammatory macrophages do not preferentially localize to the injury and instead suppress CM proliferation. Thus, pro- and anti-regenerative immune factors can now be dissected in cardioids. While we focused on the role of early, yolk sac-derived and resident-like CCR2- macrophages, it will be interesting to investigate the behavior of later, circulating monocyte-derived CCR2+ macrophages in this system.

We further demonstrate that synergistic signaling through NRG1 and FGF2 pathways is sufficient to induce trabecular CM identity and morphology in the absence of an endocardial lining, revealing that endocardial-derived NRG1 is not sufficient^60^. Thus, the absence of endocardium can be an advantage in assessing the effects and sufficiency of signaling. Future work on the full integration of circulation and endocardium will further complement the system.

HA emerged as a central mediator in establishing trabecular identity and morphology in cardioids, acting *in vivo*^8,42^ through both signaling and mechanobiological mechanisms that were previously inaccessible to experimental dissection. Although trabeculation ^6,7^ and HA^13^ were previously associated with regeneration separately in different systems, we suggest a synergistic role of trabecular signaling and HA in human fetal cardiac injury and regeneration. Consistent with this hypothesis, trabecular cardioids exhibit enhanced responsiveness to injury that is dependent on HA. This is characterized by immediate early phosphorylation of cJUN, stronger activation of TGFβ and YAP targets, and likely downstream ECM-related cytoskeletal and mechanosensing-mediated remodeling together with transient induction of CM proliferation^61–63^. These observations suggest that the developmental CM state influences injury competence and may contribute to regional differences in injury response *in vivo*, such as the heightened AP-1-mediated responsiveness of the sub-endocardial injured myocardium in patients^48^. Indeed, the reactivation of developmental mechanisms, particularly trabecular genes and morphology, is also a hallmark of dilated and hypertrophic cardiomyopathies, suggesting broader relevance in cardiac stress^64,65^.

Finally, although both YAP and WNT have been implicated in cardiac regenerative processes^20,22,36,49,66^, their specific roles in CM proliferation versus pro-regenerative remodeling can now be dissected *in vitro*. YAP is well known to convert a multitude of signaling, mechanical, and cytoskeletal inputs into coordinated organ-level transcriptional responses that control proliferation, ECM remodeling, and intercellular communication^67^, which may explain its sufficiency to induce repair, unlike WNT activation alone. In a broader context, these pro-proliferative/regenerative conditions can now be tested against anti-proliferative/regenerative factors to assess effects and potential therapies in a holistic and physiologically relevant manner. Taken together, our results indicate that specific immune recruitment, as well as coordination of both structural remodeling and proliferative signaling, are required to promote repair. This work establishes a complementary human cardiac injury *in vitro* platform for identifying and testing candidate regenerative mechanisms described in animal systems, as well as for confirming human efficacy before clinical trials.

### Limitations

The cryoinjury paradigm used here is not intended to model adult myocardial infarction but rather to reconstitute aspects of fetal-like injury responses, including macrophage subtype interactions, trabecular cardiomyocyte identity, and ECM-dependent mechanobiology involving HA. The relative immaturity of cardioids represents an inherent limitation of the system; however, modeling early developmental stages provides a rational entry point for identifying mechanisms that may underlie regenerative competence in the human heart. Progress toward modeling adult cardiac repair will require stepwise refinement of experimental systems that bridge fetal and mature tissues, while preserving key determinants of injury responsiveness. By enabling controlled dissection of how immune signals, extracellular matrix composition, and CM identity regulate remodeling and proliferative responses, this platform provides a foundation for understanding mechanisms that may ultimately support regenerative strategies in the human heart.

## Supporting information

Video S1: Anti-inflammatory macrophages migrating inside cardioid.

Video S2: Anti-inflammatory macrophage localization at injury site.

Video S3: Cardioid co-culture with different macrophage types.

Video S4: LV and TLV cardioid injury response.

## ACKNOWLEDGMENTS

We thank all laboratory members, Martina Cirigliano and Fabrizio Olmeda for their help and discussions. We are grateful to the VBC Histology & NGS, IMP/Institute for Molecular Biotechnology (IMBA) Core, and IMBA Tissue Culture Support for their services and to the Allen Institute for cell lines. We thank Life Science Editors for scientific editing. We are grateful to Elad Bassat and Miguel Torres for providing the inducible YAP construct, and to Joakim Lundeberg for supporting the spatial transcriptomics analysis. This work was funded by the Austrian Academy of Sciences (OEAW), the Era4Health Partnership CARDINNOV consortium (RECREATE) grant, and the EU Horizon Europe research and innovation program (Advanced ERC, CardioGrowth, Nr. 101201790).

## AUTHOR CONTRIBUTIONS

L.C.G. and S.M. co-designed the project and experiments and co-wrote the paper. L.C.G. developed and characterized trabecular cardioids, performed the scRNAseq experiment, and directly planned and performed the analysis with all other co-authors. T.I. established hPSC-derived macrophage generation, FUCCI and YAP5SA cell lines. L.P. performed CM proliferation and functional HA analysis. M.N. performed scRNA-seq bioinformatic analysis. A.K. and Lokesh P. established the image analysis quantification pipelines. E.L. performed VisiumHD spatial transcriptomics experiment and R.M. performed spatial bioinformatic analysis. S.H.G. generated and analyzed HREM data. K.M. performed light-sheet live imaging. All other authors performed experiments and helped with analysis. S.M. supervised the study.

## INCLUSION AND DIVERSITY

We worked to ensure diversity in experimental samples through the selection of the cell lines. While citing references scientifically relevant to this work, we also actively worked to promote gender balance in our reference list.

## METHOD DETAILS

### hPSC culture

The E8 culture system^68^ was used to cultivate all human pluripotent stem cell lines in a customized in-house medium. 0.5 percent BSA (Europa Biosciences, #EQBAH70), in-house manufactured human FGF2 (200ng/ml) or thermal stable Qkine FGF2 (#Qk053) at 5.5ng/mL), and 1.8 ng/ml TGFb1 were added to the original E8 mix (R&D RD-240-B-010). Cells were cultured on Vitronectin XF (Stem Cell Technologies) coated Eppendorf (Eppendorf SE, #0030 721.110) or TPP (TPP Techno Plastic Products AG, #92012) tissue culture-treated plates and passaged every 2-4 days at approximately 70 percent confluency using TrypLE Express Enzyme (Gibco, #12605010). The absence of Mycoplasma contamination in cells was regularly tested.

### Generation of TNNT2:FUCCI and YAP5SA hiPSCs

The TNNT2:FUCCI and YAP5SA expression cassettes were integrated into the AAVS1 locus via TALENs using AAVS1-Puro cTnT FUCCI (Addgene #136935) and YAP5SA construct (provided by Miguel Torres lab, CNIC, Spain) cloned into AAVS1-TRE3G-GFP vector (Addgene #52343). hPSCs that were approximately 70% confluent, were dissociated using StemPro™ Accutase™ Cell Dissociation Reagent (Gibco, #A1110501) and transfected using the P3 Primary Cell 4D-Nucleofector X Kit S (Lonza-BioResearch #V4XP-3032) and Amaxa 4D-Nucleofector (Lonza-BioResearch) with pulse code DS-120. Post nucleofection, cells were incubated in E8 supplemented with ROCKi (5µM) on a 6-well plate previously coated with Vitronectin XF (StemCell Technologies #7180). The following day, medium was refreshed with E8 supplemented with ROCKi. The media was replaced with E8 for the following two days. Antibiotic selection was carried out for three days with daily media changes using E8 supplemented with Puromycin (0.5µg/ml). After selection, cells were dissociated using TrypLE Express and seeded as single cells (2k cells) into a 10cm dish. Colonies were picked and transferred to a 96 well plate approximately 7 days after single cell seeding. After reaching about 70% confluency, selected colonies were dissociated and expanded to a 12 well plate for use in experiments.

### LV cardioid generation

hPSCs are seeded in a 24-well plate (TPP, #92024) at 20-30k cells per well in E8 + ROCKi (5 µM Y-27632, Tocris #1254). All differentiation media are based on CDM that consists of 5 mg/ml bovine serum albumin (Europa Biosciences, #EQBAH70) in 50% IMDM (Gibco, #21980065) plus 50% F12 NUT-MIX (Gibco, #31765068), supplemented with 1% concentrated Lipids (Gibco, #11905031), 0.004% monothioglycerol (Sigma, #M6145-100ML) and 15 mg/ml of transferrin (Roche, #10652202001). 24 hours after seeding in the 24-well plate, the cells are induced with mesoderm induction media. Mesoderm induction media is made up of CDM containing FGF2 (30 ng/ml, Cambridge University) (alternatively QKine FGF2 (5.5 ng/mL Qk053)), LY294002 (5 µM, Tocris, #1130), Activin A (5ng/mL, Cambridge University), BMP4 (10 ng/ml, R&D Systems RD-314-BP-050), and CHIR99021 (2-3 µM, R&D Systems RD-4423/50). After 36-40 hours, at d1.5, cells are dissociated with TrypLE (Gibco, #12605010) and seeded in a Corning ultra-low attachment 96 well plate (Corning, #7007) at 1500k cells/ well in Cardiac Mesoderm Patterning Media made up of CDM containing ROCKi (5 µM Y-27632, Tocris #1254), BMP4 (10ng/ml), FGF2 (8 ng/ml, Cambridge University) (alternatively, QKine FGF2 (1.466ng/mL, Qk053)), insulin (10 µg/ml), C59 (2 µM, Tocris, #5148/10) and retinoic acid (50 nM, Sigma Aldrich, #R2625). After seeding, the cells are spun down in a centrifuge for 4 mins at 140g. From d2.5 until d5.5, cells are fed every day with Cardiac Mesoderm Patterning Media without ROCKi. From d5.5 until d7.5, media is exchanged every day with Cardiomyocyte Differentiation Media CDM medium containing BMP4 (10 ng/ml), FGF2 (8 ng/ml) (alternatively, QKine FGF2 (1.466 ng/ml, Qkine053), and insulin (10 µg/ml)). This medium was termed Cardiomyocyte Specification Media. For the subsequent days of culture, media is exchanged every other day with CDM containing insulin (10 µg/ml).

### TLV cardioid generation

At day 7 of differentiation, LV cardioids are further differentiated into TLV cardioids by transferring them into CDM media containing NRG1 and FGF2 until day 10 or day 14 of differentiation (unless otherwise specified as in Fig.S2C). Media is exchanged every other day.

### hiPSC-derived macrophages generation and co-culture with cardioids

hiPSC-derived monocytes and macrophages were generated following a published protocol^31^. Monocytes were thawed into FBS-coated 6 well-plate and kept in IF9S media supplemented with M-CSF (80ng/mL, Miltenyl Biotech, #130-096-492) for 4-6days to generate M0 macrophages. M0 macrophages were then dissociated using StemPro® Accutase® Cell Dissociation Reagent (Gibco #A1110501) and seeded into 96wp at 50K cells/well in IF9S media supplemented with M-CSF. 24 hours later, polarization in pro-inflammatory and anti-inflammatory macrophages was performed by replacing the media with IF9S media containing LPS (10ng/mL, Sigma-Aldrich, #L6529) and IFN-γ (20ng/mL, PeproTech, #300-02) or IL-4 (20ng/mL, PeproTech, #200-04), respectively. At D7 of cardioid differentiation, cardioids were transferred into 96wp containing polarized macrophages and co-cultured using CDM media supplemented with 2x Insulin-Transferrin-Selenium-Ethanolamine (ITS -X) (Thermo Fisher Scientific #51500056), 64mg/L L-Ascorbic Acid 2-Phosphate (Sigma-Aldrich #A8960), 1x MEM Non-Essential Amino Acids Solution (Gibco #11140035), 1x GlutaMAX™ Supplement (ThermoFisher Scientific #35050038) and respective cytokines and cardioid treatments.

### Cryo-injury

Cardioids were temporarily transferred into a 10 cm dish without medium and placed under an EVOS microscope (Thermo Fisher). They were then contacted with a liquid N2-cooled steel rod until the wavefront of freezing tissue propagated to approximately a third of each cardioid. Cardioids were then transferred back into the 96wp for further culture and treatment.

### Enzymatic perturbation using hyaluronidase

For enzymatic perturbation cardioids were treated with hyaluronidase (1 mg/ml in the corresponding media, Sigma-Aldrich #H3506) at 37C, 5% CO2 at specified timepoints. In rescue experiments, cardioids were treated with 0.1 µg/mL to ensure optimal HAase removal upon end of treatment. Cardioids were washed 3 times with CDM media before proceeding with the rescue experiment.

### TLV cardioids rescue

Fluorinated oil (HFE-7500, fluorochem F051243) or HA (Sigma-Aldrich, cat#53747) were injected into HAase-treated TLV cardioids at D10 of differentiation. Microinjection was manually done via mouth-pipette using needles (TW100F-4, World pPrecision Instruments) pulled on a Micropipette Puller (Sutter P-97) (heat 520/pull 150/vel 100/time 150). The final volume of injected oil was adjusted to approximate the control (untreated) cardioid size. Cardioids that did not retain their oil droplet because of leakage/rupturing of the tissue were excluded from further analysis. Mock injection was performed by puncturing cardioids with a needle.

TLV cardioids were treated with HA (Sigma-Aldrich, cat#53747), added to trabeculation treatment media.

### Fixation and cryosectioning

Cardioids were fixed with 4% PFA in PBS and cryoprotected with 30% sucrose in PBS before embedding. The embedding was carried out using the O.C.T. cryo embedding medium (Scigen, #4586K1). Embedded tissues were frozen using a metal surface submerged in liquid nitrogen and stored in a 80 C freezer until sectioning on a Leica cryostat. Sections were collected on SuperFrost Plus slides (Thermo Fisher Scientific, #10149870) and kept at 20 C or 80 C until immunostaining.

### Immunostaining

To remove O.C.T., fixed specimens were washed in 1X PBS for 15 min. Optionally, tissues were placed in a permeabilization solution of 0.5% Triton-X100 (Sigma-Aldrich, #T8787) for 5 mins to increase antibody permeabilization. Tissues were then incubated in blocking solution (PBS (GIBCO, #14190094) with 4% donkey serum (Bio-Rad Laboratories, #C06SB) and 0.2% TritonX-100 for at least 30 min. Subsequently, specimens were incubated for 3 hours at room temperature or overnight at 4 C in a blocking solution containing the primary antibody. Then, a 20-minute washing in PBS with 0.1% Tween20 (Sigma-Aldrich, #P1379) was performed, followed by incubation for 1 hour at room temperature in a blocking solution containing the secondary antibody. Finally, tissues were washed in PBS with 0.1% Tween20. Slides were mounted using a fluorescence mounting medium (Dako Agilent Pathology Solutions, #S3023) and covered with a cover slip (Menzel-Gläser, #631-0853 VWR).

### In Situ Hybridization Chain reaction (HCR)

HCR fluorescent in situ was carried out using the HCR kit (v.3), purchased from Molecular Instruments (molecularinstruments.org), according to the manufacturer’s instructions with the slight modification of adding 100 mg/ml salmon sperm DNA to the pre-amplification solution and the amplification solution including the hairpins to reduce nonspecific binding. The HCR probes were designed and manufactured by Molecular Instruments.

### Scanning Electron Microscopy (SEM)

Cardioids were transferred to a mixture of 2% Paraformaldehyde (Electron Microscopy Sciences, Hatfield, PA) and 2% Glutaraldehyde (Agar Scientific, Essex, UK) in 1x PBS and incubated overnight at room temperature. Organoids were cut into halves to obtain images from the inner regions. All organoids were post-fixed in 2% osmium tetroxide (Agar Scientific, Essex, UK) for 40 min at 4°C. Samples were then 2x rinsed with PBS and 1x in ddH2O (10 min each). Chemical dehydration was performed in a 3:1 mixture of 2,2-Dimethoxypropane (DMP, Sigma- Aldrich) and ddH2O for 10 min, followed by three times anhydrous acetone and three times a 1:1 mixture of anhydrous acetone and Hexamethyldisilazane (HMDS, Sigma-Aldrich), 30 min each step. After two additional dehydration steps in pure HMDS for 1h each, samples were air dried, sputtered with a thin gold layer in a Balzers SCD 050 and examined with a Hitachi TM-1000 tabletop scanning electron microscope operated at 17 kV and equipped with a high-sensitive semiconductor BSE detector.

### High Resolution Episcopic Microscopy (HREM)

Cardioids processing and HREM data generation was performed following protocol described in Geyer et al. 2024^69^

### SimView Light-sheet live imaging

SiMView light-sheet microscope as configured in McDole, 2018^70^ with the following changes for imaging cardiods: cardioids were mounted using PEEK platforms, over which 3 mm Teflon FEP AF capillaries were placed and filled with culture media. Environmental conditions were 37C at normoxic conditions and 5% CO2.

### Image acquisition

Spinning disk confocal microscopes (Olympus spinning disk system based on an IX3 Series (IX83) inverted microscope, equipped with a Yokogawa W1 spinning disc) were used to image fixed tissue sections and whole-mount fixed cardioids. Images taken with the confocal microscope that contain more than one color are composites. Live imaging was carried out using an inverted widefield microscope for brightfield and fluorescence (Axioobserver Z1 equipped with an sCMOS camera (Hamamatsu Orca Flash 4). Cardioids in 96-well plates were also imaged using a Celigo Imaging Cytometer microscope (Nexcelom Biosciences, LLC).

### 3D Wholemount clearing and imaging of cardioids

After cardioids fixation, we performed delipidation by moving cardioids into a solution with 50% ddH2O and 25% CUBIC-L + 25% CUBIC-R1 for 3 hours at RT. Then, they are transferred into a CUBIC-L + CUBIC-R1 solution in a 1:1 ratio overnight at 37 C. The next day, cardioids are washed 3 times for 10 minutes each with PBS + 20% TWEEN20 (PBS-T). When necessary, antibody staining is performed by first leaving cardioids into blocking solution for 2 hours at RT. Primary antibodies are incubated overnight at 4C, followed by washing with PBS-T and incubation with secondary antibodies (1:250) for 2 hours at RT. Finally, cardioids are washed with PBS-T and RI matching is performed with EasyIndex 1.52 (Live Canvas Technologies, cat#EI-500-1.52) at RT for a minimum of 2 hours. After clearing, cardioids are mounted between two coverslips using ispacers (SUNJinLab, cat# IS009) and Easyindex. Cleared samples were imaged with Olympus spinning disk system equipped with a Yokogawa W1 spinning disc. Z-stacks were acquired with 3 micron z-steps.

### Flow cytometry

Cardioids (8 cardioids per condition) were dissociated using a 1 mL CM dissociation medium (Stem Cell Technologies, #05025) for 7 - 10 min at 37 C. Dissociation of CMs was stopped by adding 5 ml of the support medium. After centrifugation for 4 min at 130 g, cells were resuspended in 600 ml PBS with 0.5 mM EDTA (Biological Industries, #01-862-1B) and 10% FBS (PAA Laboratories, #A15108).

Monocytes/ macrophages were harvested at different stages of the generation for validation. Cells were filtered using a 100µm CellTrics filter and stained using fluorescent-conjugated FACS antibodies for 30min at 4 C. After staining, cells were washed once and resuspended in 500ml PBS with 10mM EDTA and 0.5% BSA.

Cells were acquired with a FACS LSR Fortessa II (BD) and analyzed with FlowJo V10 (FlowJo, LLC) software.

### RNA extraction and bulk RNA-seq preparation

RNA was isolated using an in-house RNA bead isolation kit semi-automated using KingFisher devices (KingFisher Duo Prime). Using the QuantSeq 30 mRNA-Seq Library Prep Kit FWD (Lexogen GmbH, #015), the bulk RNA-seq libraries (N=3, n=8) were generated according to the manufacturer’s instructions. After the preparation of the libraries, samples were checked for an adequate size distribution with a fragment analyzer (Advanced Analytical Technologies, Inc). Then the RNA-seq library was submitted to the Vienna Biocenter Core Facilities (VBCF) Next-Generation-Sequencing (NGS) facility for sequencing.

### Real-time quantitative polymerase chain reaction

The isolated RNA was reverse transcribed to cDNA using the Reverse Transcription Kit (Invitrogen, #18080044) with a C100 Touch Bio-Rad Thermal Cycler. Quantitative PCR was performed using the GoTaq qPCR master mix 2x (Promega, #A6001) with a Bio-Rad CFX384 Real-Time thermal cycler. Values of gene expression of each sample were obtained in triplicates. The Log-fold change of the sample from PBGD as a housekeeping gene and a pluripotent stem cell sample for normalization was calculated using a custommade script written in Python.

### Sample preparation for scRNA-seq

For scRNA-seq, 6-8 cardioids per condition and biological replicate were pooled together. We performed a scRNA-seq time-course analysis on LV and TLV cardioids cultured alone or with anti-inflammatory macrophages (aiMPs), in the presence and absence of cryo-injury, at 1 and 3 days post-injury (dpi). Two biological replicates were used per condition. Each biological replicate is composed of two independent scale bio experiments (full plate and extension plate). Cardioids were dissociated using 1 mL CM dissociation medium (Stem Cell Technologies, #05025) for 7-10 min at 37 C. Dissociation of CMs was stopped by adding 5 ml of the support medium. The cell suspension was spun down at 400 g for 4min at 4 C. The supernatant was aspirated, and the cell pellet was resuspended in 400ml ice-cold PBS/BSA (1%). Single cell suspensions of cardioids were submitted to the VBCF NGS facility (Vienna, Austria) for fixation and library preparation. Cell fixation followed the ScaleBio ‘Low Volume Fixation’ User Demonstrated Protocol (Document 1020807) using the ‘ScaleBio Sample Fixation Kit’ (PN2020001, Scale Biosciences, San Diego, CA). Single cell cDNA libraries were prepared using the ScaleBio ‘Single Cell RNA Sequencing v1.1’ and ‘Single Cell RNA Extended Throughput v1.1’ kits (PN950884 and PN936360, Scale Biosciences, San Diego, CA).

### Spatial data generation

Four cardioids per experimental group were embedded in OCT (Scigen, #4586K1) medium and snap-frozen using a metal surface submerged in liquid nitrogen. 10 µm thick cryosections were collected onto Superfrost Plus glass slides (Thermo-Fisher, 22-037-246) across the entire depth of the cardioids. Following 10 min fixation in 4% formaldehyde (Thermo-Fisher, 28908), hematoxylin-eosin staining (3 min in Gill No. 2 hematoxylin, Sigma-Aldrich, GHS232; 45 sec in aqueous eosin Y, Sigma-Aldrich, HZ110216, diluted 1:10 in Tris-acetic acid buffer pH 6.0) was performed on evenly paced sections to determine the optimal depth for subsequent spatial transcriptomics analysis, based on the size of the cryoinjured region captured within the sections. Spatial gene expression libraries were prepared from a selected section of each sample using the Visium HD Gene Expression kit according to the manufacturer’s instructions (10× Genomics, CG000685 User Guide, Rev C). Prior to the Visium workflow, tissues were fixed, baked, and subjected to hematoxylin-eosin staining following the RNA Rescue Spatial Transcriptomics (RRST) protocol^71^. Tissue images were taken at ×20 magnification using the Metafer Slide Scanning platform (microscope, AxioImager.Z2 with ScopeLED Illumination, Zeiss; camera, CoolCube 4 m, MetaSystems; objective, Plan-Apochromat ×20/0.80 M27, Zeiss; software, Metafer5). Raw images were stitched with VSlide software (MetaSystems). In total, 4 Visium libraries were prepared from two biological replicates (#83, #96) of the LV and TLV (marked by NF in the bioinformatic pipeline) models. Libraries were sequenced by using the Illumina NextSeq 2000 platform, in which the length of read 1 was 43 bp and the length of read 2 was 50 bp

### Contraction Analysis

Cardioids were fed fresh CDMI media 1-2 hours before recordings. The 96-well plate was placed in an environmentally controlled stage incubator (37◦C, 5% CO2, water-saturated air atmosphere, Okolab Inc, Burlingame, CA, USA). Each well was imaged using widefield phase-contrast microscopy (Axioobserver Z1 (inverted) with sCMOS camera, Zeis) at 100 frames per second for 30 seconds. Videos were then analyzed using MUSCLEMOTION; the data was read into custom-made software for reported calculations. Percent beating was defined by whether the cardioid beat once within the entirety of the recording. Beats per minute were calculated by counting the total number of beats in the video, dividing them by the length of the video in seconds, and multiplying by 60.

## QUANTIFICATION AND STATISTICAL ANALYSIS

### Measuring Nuclear Densities

Images of cardioids were analyzed to quantify the spatial density of nuclei that are positive for cardiomyocyte signal. Cardioids were cut in sections and then stained for nuclei (DAPI) and cardiomyocytes (TNNT2). Maximum intensity projections were generated from 3D image stacks to collapse the signal into a 2D representation. Binary masks for nuclei and cardiomyocytes were created using a fixed intensity threshold for each channel. The cardiomyocyte mask was dilated to account for spatial spread, and nuclei were counted only within regions positive for cardiomyocyte signal by multiplying the respective binary masks. To quantify local nuclear density, the resulting binary mask for nuclei (restricted to cardiomyocyte-positive regions) was divided into non-overlapping square bins of fixed size (100 x 100 pixels). Within each bin, the density was calculated as the fraction of pixels occupied by nuclei:

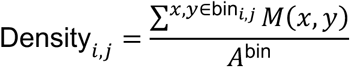

where *M*(*x*, *y*) is the binary mask value (1 for nuclear-positive and cardiomyocyte-positive, 0 otherwise), and *A*^bin^ is the total number of pixels in the bin (10^’^). Thus, the density represents the local proportion of cardiomyocyte-associated nuclear signal within each spatial region. The distribution of density values across bins was visualized as a heatmap. The density values for bins containing exclusively the cavity or endodermal tissue would be 0. While summarizing the mean, median, and standard deviation of densities, bins with 0 density were excluded. Thus, nuclear densities were measured only in tissue regions. All image processing steps, including segmentation, masking, binning, and quantification, were automated using Python scripts. Quantitative results, including total nuclear area, summed and mean intensities, and density statistics, were saved in structured JSON and CSV formats for downstream analysis.

### Visualizing Multi-channel images

For visualization of multichannel images, each wavelength was processed and displayed as a separate subplot within a montage. Prior to plotting, intensity normalization was performed on each wavelength by scaling pixel values between user-defined lower and upper bounds, as specified in the metadata file. Specifically, each pixel intensity (I) was normalized using the formula:

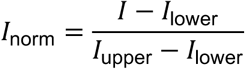

where *I*_lower_ and *I*_upper_ are the channel-specific lower and upper intensity bounds, respectively. Values exceeding the upper bound were capped at 1, and those below the lower bound were set to zero, ensuring consistent dynamic range across channels. Custom colormaps were applied to each normalized channel to facilitate visual discrimination of different wavelengths. The resulting subplots enable qualitative comparison of signal distribution and intensity across all channels within the same field of view.

### Wholemount image analysis (PHH3^+^ nuclei quantification)

Cardioids were stained for proliferating nuclei (PHH3^+^) and imaged as z-stacks with a step size of 3 µm. All image stacks were resized to 512 × 512 pixels and standardized to 100 z-slices to ensure consistency across samples. Segmentation of pH3-positive nuclei was performed using a U-Net–based deep learning model trained on 2D z-slices. Ground truth masks were generated by applying Triangle thresholding to 1,100 annotated 2D images of pH3 staining. The trained model was then used to predict segmentation masks for each slice. Counting of nuclei was performed using connected component labeling (via the scikit-image label function) across the segmentation masks in 3D.

### Cardioid size analysis

Cardioid area was determined from widefield images using the MOrgAna tool for quantitative analysis of organoids^72^.

### Bulk RNA-seq analysis

Bulk RNA-seq samples displayed in Fig. 2 was generated using Lexogen’s Quantseq. Initial read processing was performed by removing adapter, polyA and low-quality bases from the 3’ end (ref=polyA.fa.gz,truseq.fa.gz k=13 ktrim=r useshortkmers=t mink=5 qtrim=r trimq=10 minlength=20) using BBDuk v38.06 and filtering for abundant sequences included in the iGenomes UCSC hg38 reference (human rDNA, human mitochondrial chromosome, phiX174 genome, adapter) using bowtie2 v2.3.4.1. Remaining reads were analyzed using genome and gene annotation for the GRCh38/hg38 assembly obtained from Homo sapiens Ensembl release 94. Reads were aligned to the genome using star v2.6.0c and reads in genes were counted with featureCounts (subread v1.6.2) with strand-specific read counting for QuantSeq experiments (-s 1).

Bulk RNA-seq samples generated with the Lexogen’s Quantseq kit with UMI extension (in Fig. 3) were preprocessed using umi2index (Lexogen) to add the UMI sequence to the read identifier, and trimmed using BBDuk v38.06 (ref=polyA.fa.gz,truseq.fa.gz k=13 ktrim=r useshortkmers=t mink=5 qtrim=r trimq=10 minlength=20). Reads mapping to abundant sequences included in the iGenomes NCBI GRCh38 references were removed using bowtie2 v2.3.4.1 alignment. The remaining reads were analyzed using genome and gene annotation for the GRCh38 assembly obtained from Homo sapiens Ensembl release 112. Reads were aligned to the genome using star v2.6.0c, and reads in genes were counted with featureCounts (subread v1.6.2) using strand-specific read counting (-s 1). Differential gene expression analysis on counts was performed using DESeq2 v1.18.1; in Fig. 3 factors of unwanted variation estimated using RUVSeq v1.40.0 (RUVr, k=2) were removed, independently for TLV and LV samples.

### Single-cell RNA-seq analysis

Raw ScaleBio reads were processed using the ScaleBio ScaleRna Single-cell RNA Nextflow workflow v1.6.3 with the vendor-provided library configuration file (libV1.1.json) and genome annotation (Homo_sapiens.GRCh38.103.biotypeFiltR.gtf). Further processing of the scRNAseq data was performed in R software v4.4.2 with Seurat v5.1.0. Adaptive per sample cell filtering was carried out using scDblFinder providing expected doublet rate of 0.047, and applying a lower gene count threshold (>2 MADs). Further single-cell data processing of cell subsets after low-quality cell removal was following the same processing pipeline: log-normalization and scaling, dimensionality reduction using PCA on the top 2000 most variable genes, batch correction across replicates and plates with Harmony v1.2.3 applied to the first 20 principal components, UMAP visualization using the top 20 harmony embeddings. Gene set activity on a single cell level was computed using the AddModuleScore function. Activities of core pathways were inferred using decoupleR v2.12.0 applying PROGENy model weights of the top 500 pathway-responsive genes and the Multivariate Linear Model (MLM) approach, where positive scores (t-values) reflect increased pathway activity. Mean activities per group across pathways were visualized using pheatmap. Differential expression testing comparing conditions within broad celltypes was performed using MAST restricted to genes expressed in at least 1% of cells, including the replicate as a latent variable and run with both uncorrected STARsolo counts as well as cellbender v0.3.0 denoised counts as input. Genes were classified as differentially expressed (DEGs) if they met the significance threshold (adjusted p-value < 0.01) and were expressed in >5% of cells in the condition with higher mean expression for both uncorrected and denoised counts. In case of the M2 effect comparisons a log-fold change threshold of 0.2 was applied in addition to limit the number of DEGs. For downstream biological interpretation, DEG functional enrichment was assessed with clusterProfiler and the BioPlanet_2019 pathway database. IPA Upstream Regulatory Analysis was conducted for the identification of upstream regulators of DEGs.Single-cell enrichment analysis of the BioPlanet_2019 collection of biological pathways and the complete IPA-derived YAP upregulated target list were performed performed in python with decoupler v2.1.1, the univariate linear model (ulm) method, and plotting a heatmap of the mean expression values per group using scanpy.pl.matrixplot.

### Spatial data processing

The sequenced libraries were processed with Space Ranger (v 3.1.3; 10x Genomics) using the built-in human reference genome (GRCh38-2020-A transcriptome, Ensembl 98) as alignment reference. The remaining Visium HD bins not covering the tissue sections after Space Ranger were manually filtered using Loupe Browser (v. 8.0.0; 10x Genomics).

The down-processing and data analyses were performed in R Statistical Software (v4.4.3; R Core Team 2021) using Seurat (v5.3.0) and semla (v1.3.1). Spatial objects for all sections were created using the ReadVisiumData function from semla. The Visium HD resolutions of 8 µm and 16 µm were used and described as:

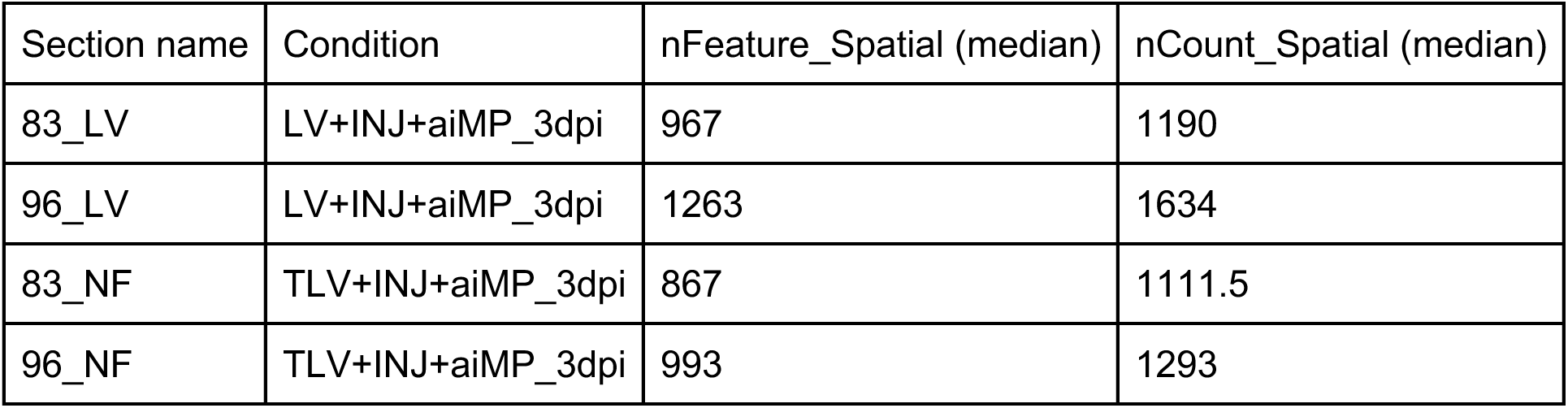

Each tissue section was then normalized and scaled individually using the NormalizeData and ScaleData functions from the Seurat package with default parameters, respectively. The most variable genes were computed using the FindVariableFeatures function from Seurat, targeting the first 3,000 most variable genes. Then, PCA dimensionality reduction was performed with the RunPCA function from Seurat with default parameters prior to the shared-nearest-neighbor (SNN) graph construction using the FindNeighbors function from Seurat with the first 30 PCs. Louvain clustering was then performed on the neighbor’s graph at multiple resolutions, easing exploration. Specifically, at 8 µm Visium HD bin resolution, clustering was performed at resolutions of 0.2, 0.4, 0.6, 0.8, and 1.0. For 16 µm Visium HD bin resolution, clustering was performed at resolutions of 0.2, 0.4, 0.6, 1.0, and 1.2. The clusters at all resolutions were mapped to the spatial data and visualized using the MapLabels function from the semla package. To understand the evolution and distribution of clusters at increasing resolution, we used the Clustree package (v0.5.1). The FindAllMarkers (logfc.threshold = 0.5, only.pos = TRUE, min.pct = 0.05, max.cells.per.ident = 300) function from Seurat was applied to identify differentially expressed genes (DEG) in each section at all the specified clustering resolutions. The FindAllMarkers function employs the two-sided Wilcoxon Rank Sum Test and accounts for multiple testing by incorporating the Bonferroni correction across all genes in the data. Individual spatial gene plotting was performed on the normalized data assay using the MapFeatures function from semla. We also performed enrichment of specific gene sets using the AddModuleScore function with default parameters from Seurat.

### Spatial mapping by Cell type deconvolution

Cell type mapping was computed separately for the two LV and two TLV sections with the corresponding single-cell datasets. The deconvolution of the single-cell clusters onto the spatial data was performed using the non-negative least squares (NNLS) method from the RcppML package (v0.3.7) implemented in semla with the RunNNLS function. NNLS successfully decomposes the spatial spot data (at 8 and 16 µm resolution) in estimating cell type proportions from the single-cell clusters used as input. The top 5,000 most variable features were computed using the FindVariableFeatures function from Seurat and utilized as the single-cell input, along with the processed Visium HD section. Each cell type population was visualized spatially using the MapFeatures function from semla, with the max_cutoff parameter set to 0.99 to exclude the top 1 percent of deconvolution values, thereby mitigating the effect of extreme proportions for visualization purposes.

### Statistics

Information on the statistics used in the experiments can be found in the Figure Legends, such as N (biological replicates, performed with different cell batches of different passages), n (technical replicates, performed with the same cell batch of the same passage) numbers, statistical test used, and variation measure used. We ran normality tests and outlier tests on all datasets before running the one-way ANOVA assuming normality. Other statistical details are described in the respective Quantification Methods sections.

## SUPPLEMENTAL INFORMATION

### SUPPLEMENTARY FIGURE LEGENDS

**Figure S1:**
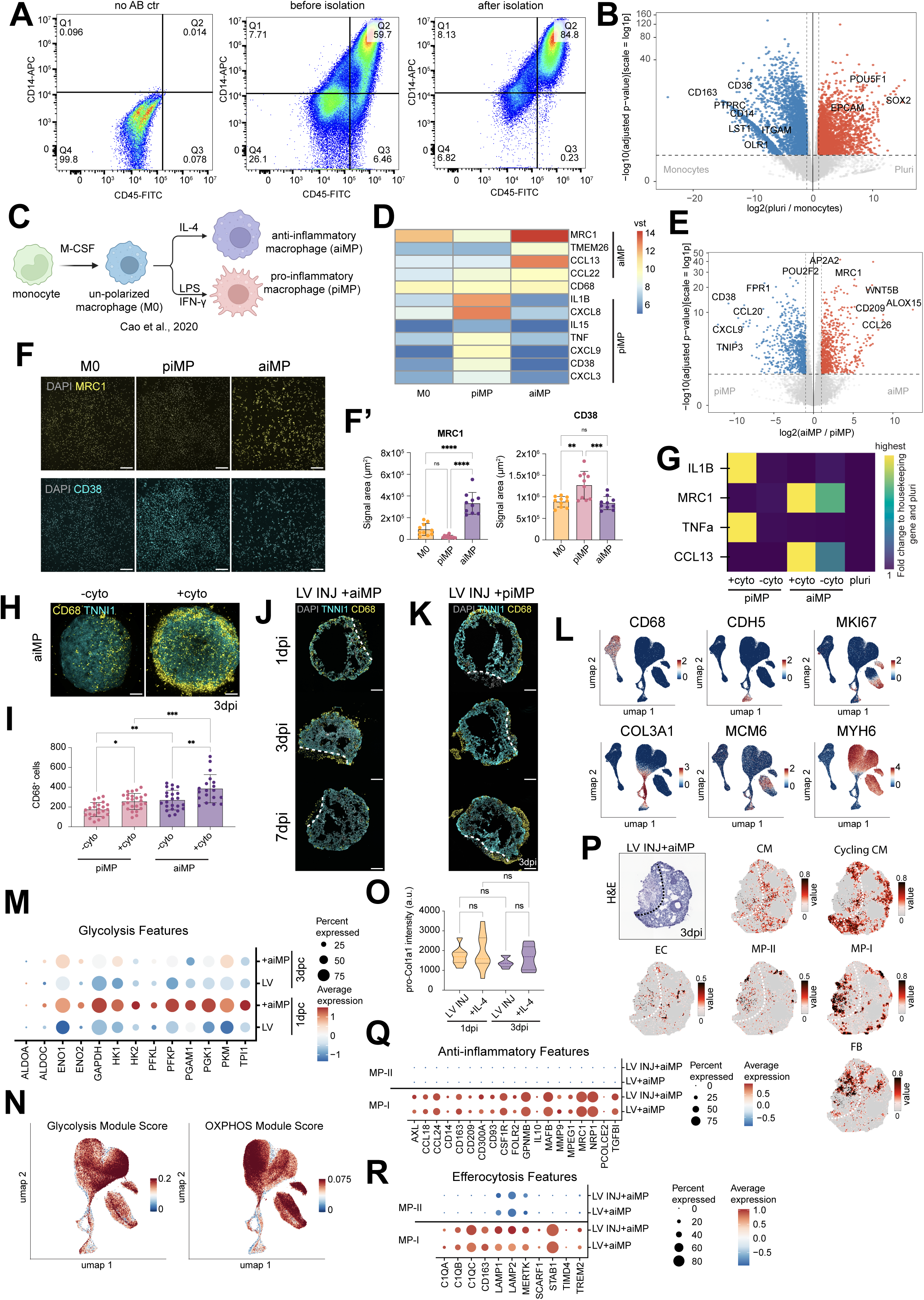
Macrophages migrate to the injury site and contribute to ECM remodeling, related to Figure 1. **A.** Representative flow cytometry plots of WTC-hPSC-derived monocytes **B and E.** RNA-seq volcano plot of differentially expressed genes in indicated conditions **C.** Schematic protocol of hPSC-derived macrophage polarization **D.** RNA-seq expression heatmap of anti-inflammatory and pro-inflammatory marker genes in indicated conditions **F.** MRC1 immunostaining and CD38 HCR staining of unpolarized (M0), pro-inflammatory (piMPs) and anti-inflammatory (aiMPs) macrophages and respective quantification (F’) (N=2, n=5) **G.** RT-qPCR expression heatmap of pro- and anti-inflammatory marker genes in indicated conditions **H.** Representative whole-mount images stained for CD68 (yellow) of LV cardioids co-cultured with aiMPs untreated and treated with respective polarizing factors (cyto). Polarizing factor is IL-4 in co-culture with aiMPs, and IFN-γ and LPS in co-culture with piMPs. **I**. Quantification of CD68+ cells in specified conditions (N=3, n=5-8) **J.** Representative cryo-sections of injured LV cardioids co-cultured with aiMPs immunostained for CD68 at 1dpi, 3dpi and 7dpi **K.** Representative cryo-sections of injured LV cardioids co-cultured with piMPs immunostained for CD68 at 3dpi **L.** Expression of marker genes for each cell types cluster plotted on UMAP **M.** Dotplot showing expression of glycolytic enzymes in cardiomyocytes in indicated single cell RNA-seq conditions **N.** Expression of glycolysis and OXPHOS gene modules **O.** Quantification of pro-Col1A1 signal intensity in the injury site from whole-mount images of cryo-injured LV cardioids with or without polarizing factor (IL-4) (N=2, n=3-5) **P.** Dotplot showing expression of anti-inflammatory markers in MP-I and MP-II cell types in indicated single cell RNA-seq conditions **Q.** Dotplot showing expression of efferocytosis-associated in MP-I and MP-II cell types in indicated single cell RNA-seq conditions **R.** Spatial mapping of scRNA-seq cell type clusters on sections of injured LV cardioid co-cultured with aiMPs at 3dpi. Scalebar: cell type proportion. Scalebar is 200 µm. Dashed line highlights the injured area. Bar graphs show mean ± SD. Statistics: one-way ANOVA. *p < 0.05, **p < 0.01, ***p < 0.001, ****p < 0.0001.

**Figure S2:**
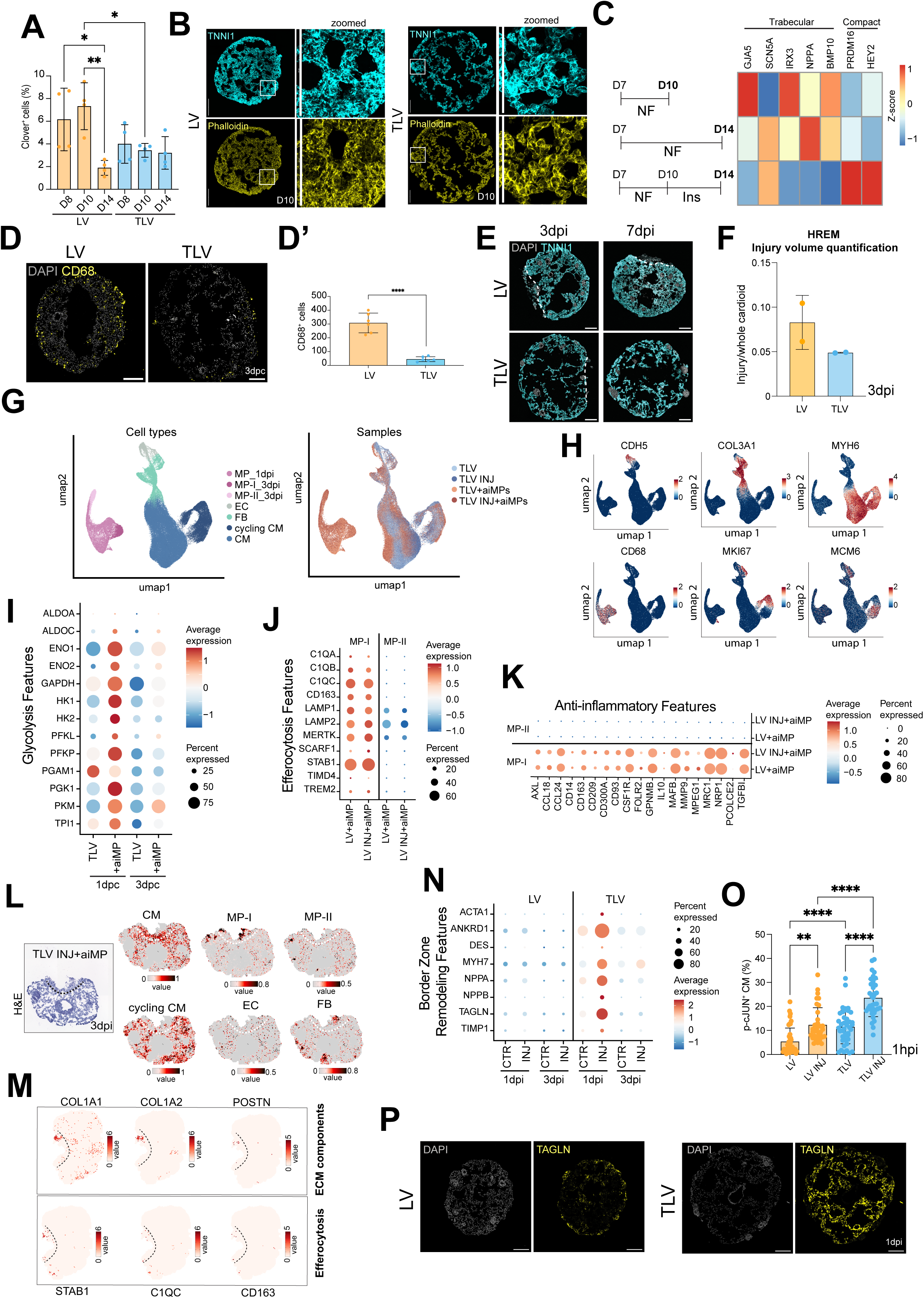
Synergistic NRG1 and FGF2 signaling induces trabecular identity, morphology, and injury remodeling, related to Figure 2. **A.** Time course quantification of Clover+ cells in TNNT2:FUCCI-hPSC-derived LV and TLV cardioids (N=4, n=8) **B.** TNNI1 and Phalloidin staining of LV and TLV cardioid cryosections at D10 **C.** RNA-seq expression heatmap of trabecular and compact marker genes in indicated condition. Harvesting timepoint is indicated in bold. **D.** CD68 immunostaining in LV and TLV cardioid cryosections at 3 days post co-culture (3dpc) and quantification (D’) (N=2, n=3), Statistics: Student’s t-test **E.** Representative cryosection of TNNI1:GFP-hPSC-derived cryoinjured LV and TLV cardioids at 3 and 7dpi **F.** Injury volume quantification from HREM imaging of cryo-injured LV and TLV cardioids at 3dpi; Injury volume is normalized to the whole cardioid volume (N=2; n=1). **G.** UMAP representation of scRNA-seq data showing distinct groups organised by cell type or sample origin **H.** Expression of marker genes for each cell type cluster **I.** Dotplot showing expression of glycolytic enzymes in indicated single cell RNA-seq conditions **J.** Dotplot showing expression of genes belonging to “Muscle contraction” Bioplanet term in cardiomyocytes across indicated conditions **K.** Spatial mapping of scRNA-seq cell type clusters (from UMAP in Fig. S2G) on sections of injured TLV cardioid co-cultured with aiMPs at 3dpi. Scalebar: cell type proportion. **L.** Spatial features plot of efferocytosis and ECM-remodelling genes in cryoinjured TLV cardioids co-cultured with aiMPs at 3dpi. Scalebar: normalized expression. **M.** phospho-cJUN+ CM quantification in indicated conditions at 1 hour post injury (hpi). **N.** TAGLN immunostaining in un-injured LV and TLV cardioids at 1dpi. Scalebar is 200 µm. Dashed line highlights the injured area. Bar graphs show mean ± SD. Statistics: One-way ANOVA, except where specified. *p < 0.05, **p < 0.01, ***p < 0.001, ****p < 0.0001.

**Figure S3:**
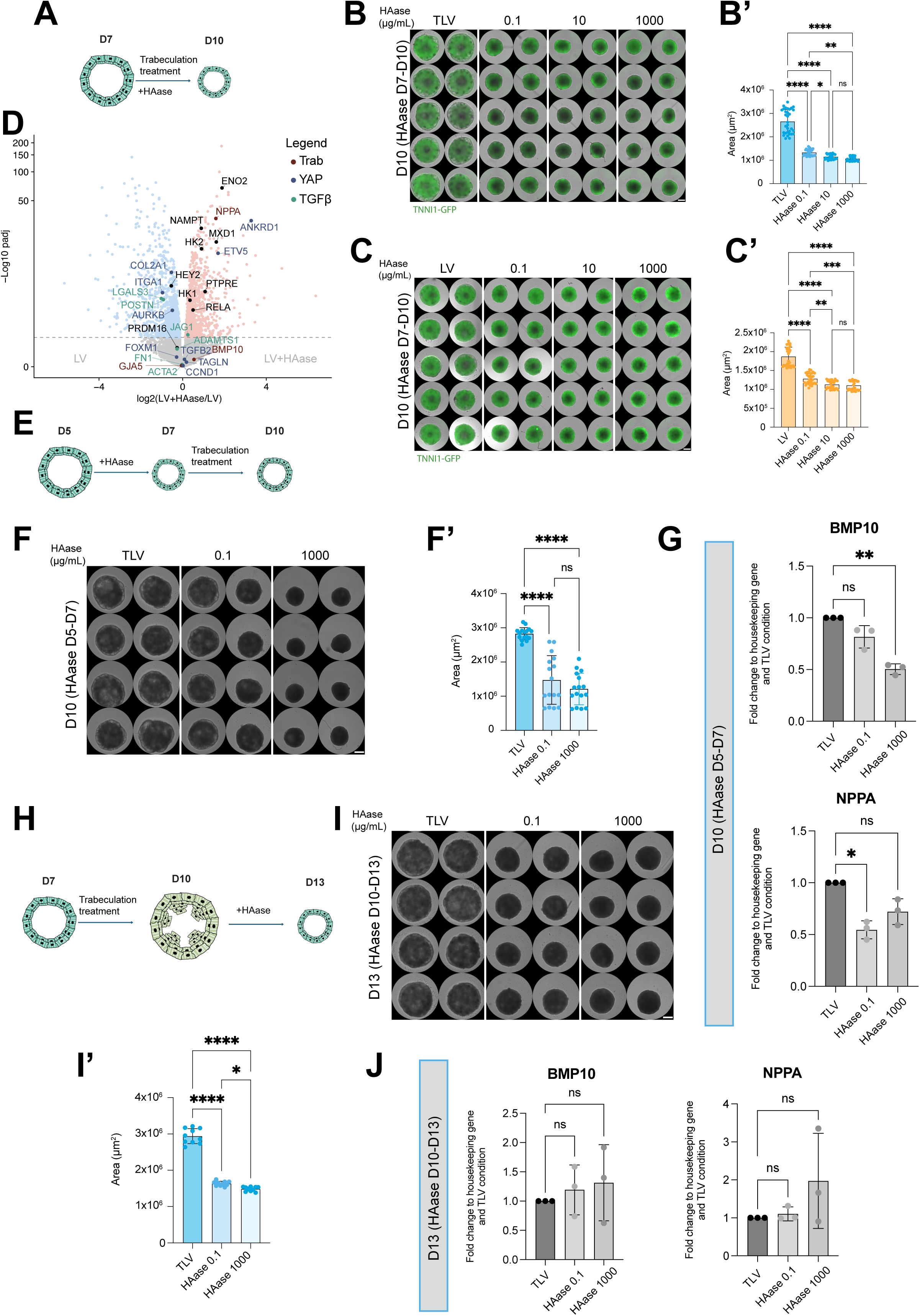
Hyaluronic acid regulates trabecular identity and injury remodeling, related to Figure 3. **A, E and H.** Schematic protocol representation of cardioids treated with HAase and/or trabeculation treatment at indicated timepoints **B.** Widefield images of TNN1:GFP-hPSC-derived TLV cardioids at D10, treated with indicated concentrations of HAase from D7 to D10, with size quantification (B’) (N=3, n=8-12) **C.** Widefield images of TNN1:GFP-hPSC-derived LV cardioids at D10, treated with indicated concentrations of HAase from D7 to D10, with size quantification (C’) (N=3, n=7-12) **D.** RNA-seq volcano plot showing differentially expressed genes in indicated conditions. **F.** Widefield images of WT-hPSC-derived TLV cardioids at D10, treated with indicated concentrations of HAase from D5 to D7, with size quantification (F’) (N=2, n=8 ) **G.** RT-qPCR of NPPA and BMP10 at D10 in conditions from panel E (N=3, n=8) **I.** Widefield images of WT-hPSC-derived TLV cardioids at D13, treated with indicated concentrations of HAase from D10 to D13, with size quantification (I’) (N=2, n=8 ) **J.** RT-qPCR of NPPA and BMP10 at D13 in conditions from panel H (N=3, n=8) Scalebar is 500 µm. Bar graphs show mean ± SD. Statistics: one-way ANOVA. *p < 0.05, **p < 0.01, ***p < 0.001, ****p < 0.0001.

**Figure S4:**
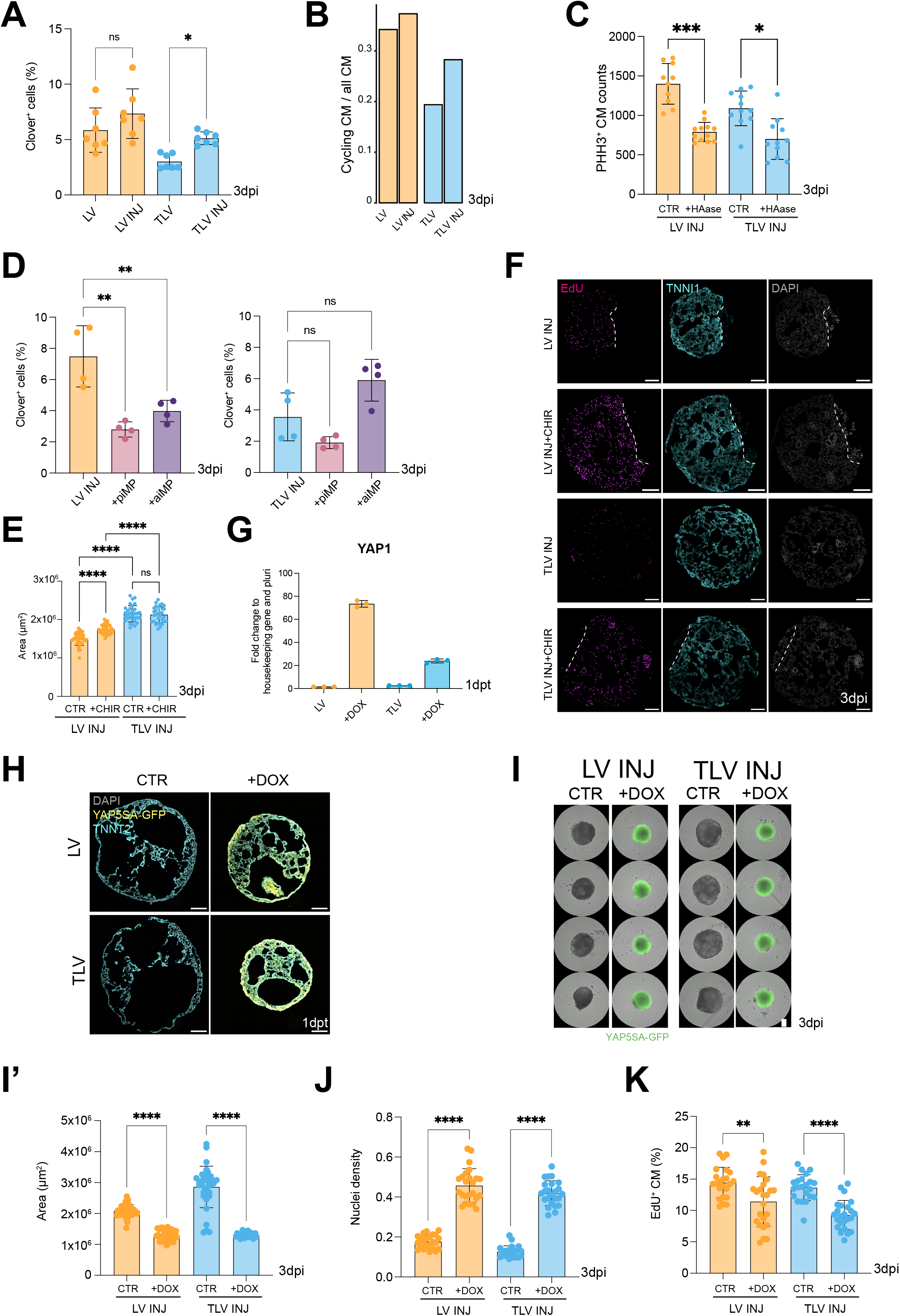
Integrated cardiomyocyte proliferation and remodeling is necessary for cardioid injury repair, related to Figure 4. **A and D.** Clover+ cells quantification from TNNT2:FUCCI hPSC-derived LV and TLV cardioids in specified conditions at 3dpi (A: N=7, n=8) (D: N=4, n=8). Statistics: Student’s t-test **B.** Ratio of cycling CM over all CM in indicated conditions from single cell RNA-seq data **C.** Quantification of PHH3+ CM from whole-mount images of indicated conditions at 3dpi (N=3, n=3-4). **E**. Size quantification of indicated conditions at 3dpi (N=4, n=8) **F.** Representative cryo-sections of injured LV and TLV cardioids labeled for EdU in indicated conditions at 3dpi **G.** RT-qPCR of YAP1 in indicated conditions at 1dpt (1 day post treatment) **H.** Representative cryo-sections of untreated and DOX-treated YAP5SA:GFP-hPSC-derived LV cardioid at 1dpt **I.** Widefield images of YAP5SA:GFP-hPSC-derived LV and TLV cardioids with size quantification (I’) of specified treatments at 3dpi (N=3, n=12). Scale bar is 500 µm **J.** Nuclei density quantification of indicated conditions at 3dpi **K.** Quantification of EdU+ CMs in indicated conditions at 3dpi. Scalebar is 200 µm, except where specified. Bar graphs show mean ± SD. Statistics: one-way ANOVA, except where specified. *p < 0.05, **p < 0.01,***p < 0.001, ****p < 0.0001.

## SUPPLEMENTARY VIDEO LEGENDS

Video S1: **Anti-inflammatory macrophages migrating inside cardioid.** Time-lapse (2 days – 3 days post injury (dpi)) SimView light sheet live imaging of LV cardioid co-cultured with anti-inflammatory macrophages (aiMP). Panels show two camera angles of the same cardioid. Green: TNNI1, Red: aiMP

Video S2: **Anti-inflammatory macrophage localization at injury site.** Time-lapse (5dpi - 6dpi) SimView light sheet microscopy imaging of cryo-injured LV cardioid co-cultured with anti-inflammatory macrophages (aiMP). Blue: nuclear dye, Green: TNNI1, Red: aiMP

Video S3: **Cardioid co-culture with different macrophage types.** Time-lapse (1dpi - 7dpi) imaging of cardioids co-cultured with hPSC-derived macrophages. Left panel: pro-inflammatory macrophage (piMP) co-culture. Right panel: anti-inflammatory macrophage (aiMP) co-culture. Scale bar: 500µm

Video S4: **LV and TLV cardioid injury response.** Time-lapse (1dpi - 7dpi) imaging of cryo-injured LV (left panel) and TLV (right panel) cardioid. Scale bar: 500µm

